## Supplementary Information for "Inferring kinetic rate constants from single-molecule FRET trajectories – a blind benchmark of kinetic analysis tools"

Götz et al.

**1 Supplementary Figures** 2

Supplementary Figure 1: Equilibrium constants of the kinetics between two states shown in Figure 2. 2
Supplementary Figure 2: Supplementary results from experimental data with high sampling rate and low SNR. 3
Supplementary Figure 3: Validation of the simulated datasets. 4
Supplementary Figure 4: Quantitative comparison of the four most accurately inferred models shown in Fig. 4. 5
Supplementary Figure 5: FRET efficiency histograms and all inferred FRET states for the experimental datasets shown in Figure 5. 6
Supplementary Figure 6: Comparison of the kinetic models with three FRET states inferred for the datasets shown in Figure 5. 7
Supplementary Figure 7: Comparison of the kinetic models with four FRET states inferred for the datasets shown in Figure 5 8

**2 Supplementary Notes** 9

Supplementary Note 1: Simulation of smFRET trajectories 9
Supplementary Note 2: Estimated minimal uncertainty of rate constants inferred from simulations 11
Supplementary Note 3: Simulation of cumulative dwell-time distributions from inferred kinetic models 11
Supplementary Note 4: A simple file format for smFRET trajectories 12

**3 Supplementary Methods** (numbered as in the main text) 13
Supplementary Method 1: Pomegranate 13
Supplementary Method 2: Tracy 15
Supplementary Method 3: FRETboard 16
Supplementary Method 4: Hidden-Markury 18
Supplementary Method 5 & 6: SMACKS 20
Supplementary Method 7: Correlation 22
Supplementary Method 8: Edge finding (CK) 30
Supplementary Method 9: Edge finding (k-means) 31
Supplementary Method 10: Step finding 33
Supplementary Method 11: STaSI 36
Supplementary Method 12 & 13: MASH-FRET (bootstrap & probabilistic) 38
Supplementary Method 14: postFRET 43

**4 Supplementary Tables** 49

Supplementary Tables 1: Inferred kinetic models for the data shown in Fig. 4. 49
Supplementary Tables 2: Inferred kinetic models for the data shown in Fig. 5a-c. 52
Supplementary Tables 3: Inferred kinetic models for the data shown in Fig. 5d-f. 55
Supplementary Tables 4: Inferred kinetic models for the data shown in Fig. 5g-i. 58

**5 Supplementary References** 61

### 1 Supplementary Figures

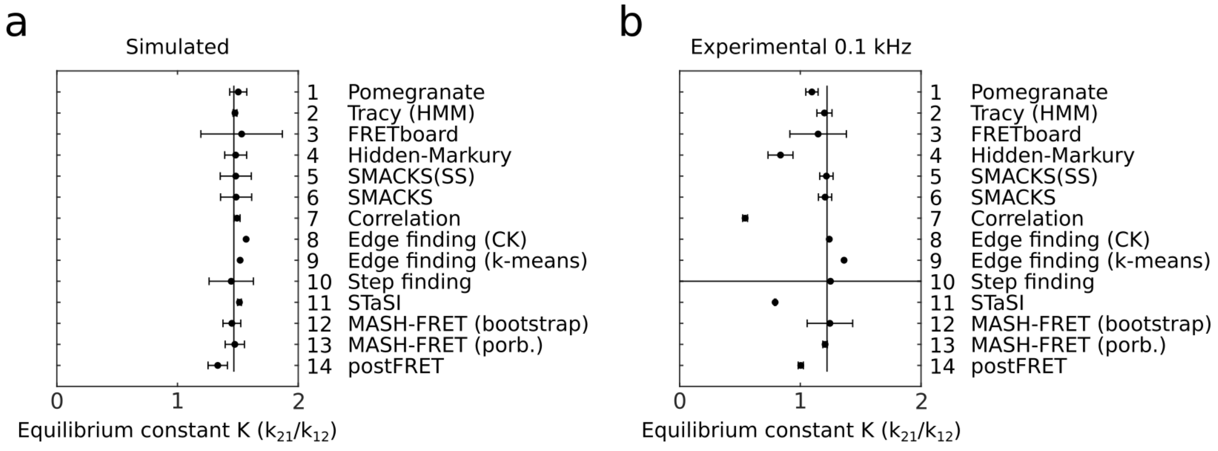


**Supplementary Figure 1 | Equilibrium constants of the kinetics between two states shown in Figure 2. a** The inferred equilibrium constant for the simulated dataset. The vertical line indicates the ground truth value. **b** The inferred equilibrium constant for the experimental data with 0.1 kHz sampling rate. The vertical black line indicates the ratio of the two well-separated FRET efficiency populations, as estimated by dividing the number of datapoints with FRET E < 0.5 by those with FRET E > 0.5.

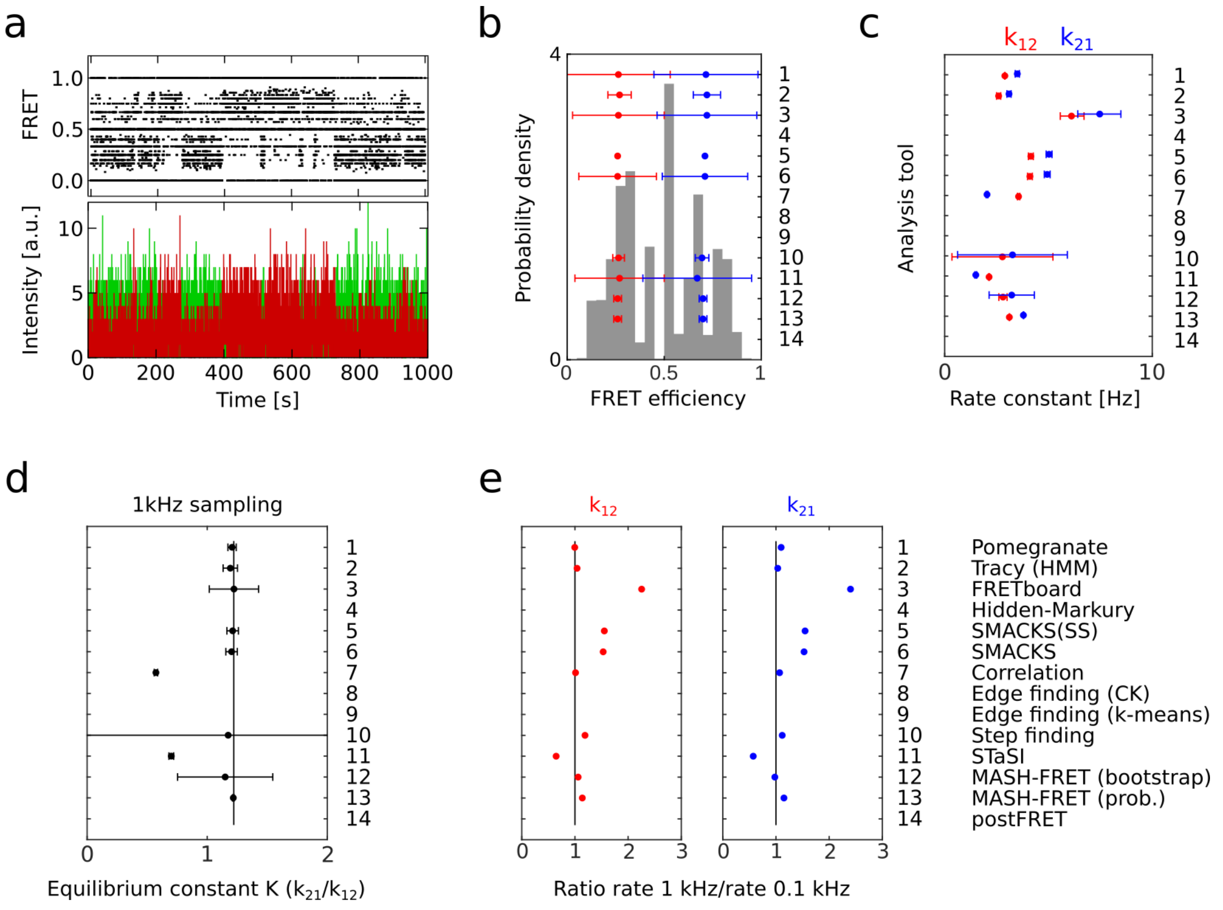


**Supplementary Figure 2 | Supplementary results from experimental data with high sampling rate and low SNR. a** Donor (green) and acceptor (red) fluorescence and FRET efficiency (FRET, black) trace for the molecule shown in Fig. 2e, with 1 kHz sampling rate. **b** Corresponding FRET efficiency histogram (gray) and inferred FRET efficiencies in red and blue. **c** Inferred rate constants from experimental data (using 1 ms time bins resulting in 1 kHz sampling). **d** Equilibrium constant for the experimental datasets with 1 kHz sampling. The vertical black line indicates the population ratio as estimated from the FRET efficiency histogram at 0.1 kHz sampling (Fig. 2f) by dividing the number of observations with FRET $E < 0.5$ by those with FRET $E > 0.5$. **e** Ratio of the rate constant inferred from data with 1 kHz vs 0.1 kHz sampling for rate $k_{12}$ (red) and $k_{21}$ (blue). The black line indicates a ratio of one, i.e., rate constants inferred for both sampling rates are equal. A ratio above one means that the rate constant inferred from the 1 kHz dataset is larger than the one inferred from the 0.1 kHz dataset.

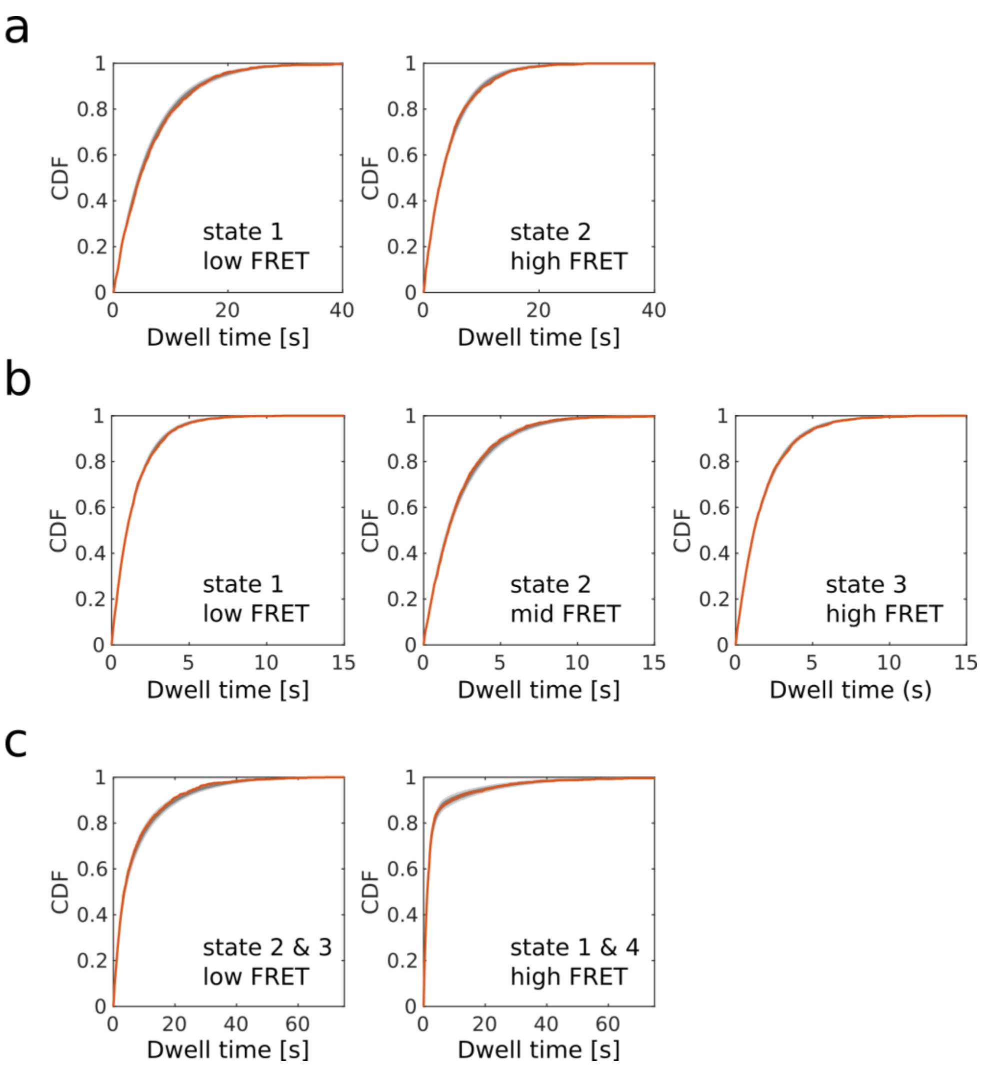


**Supplementary Figure 3 | Validation of the simulated datasets** using cumulative distribution functions (CDF) of the dwell times. The orange line represents the simulated data used in this study. The spread between 500 datasets obtained from simulations with identical parameters is shown in dark and light grey intervals representing one and two standard deviations around the mean, respectively. **a** For the simulated data shown in Fig. 2. **b** For the simulated data shown in Fig. 3. **c** For the simulated data shown in Fig. 4.

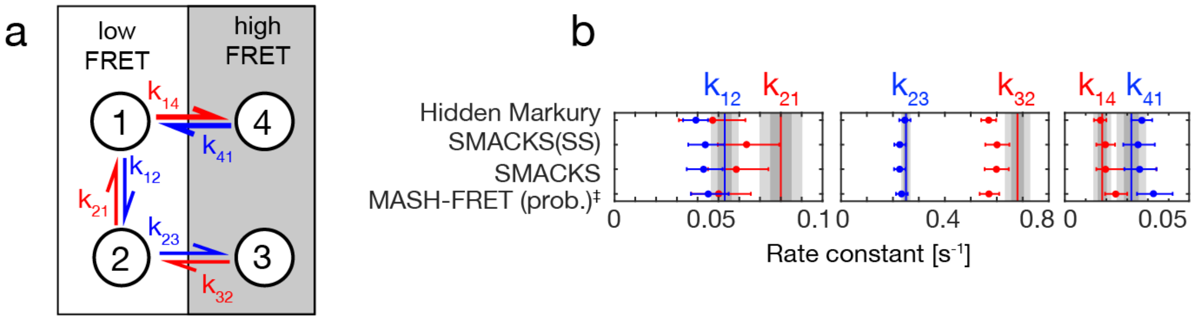


**Supplementary Figure 4 | Quantitative comparison of the four most accurately inferred models shown in Fig. 4. a** Illustration of the simulated kinetic model for reference (same as Fig. 4a). **b** The kinetic rate constants: the GT values are represented as red and blue vertical lines. The intrinsic uncertainty of the dataset is shown as dark gray (1σ) and light gray (2σ) intervals. Beyond the six displayed rate constants, these additional rate constants were inferred: for Hidden Markury $k_{31}$ = 0.045 and $k_{34}$ = 0.003, for SMACKS $k_{13}$ = 0.0001, $k_{31}$ = 0.0055, $k_{34}$ = 0.0034, for MASH-FRET (prob.) $k_{31}$ = 0.033. All inferred rate constants by all tools are reported in the Supplementary Tables 1 and in the Supplementary Datafiles. ‡ indicates values provided after the GT was known.

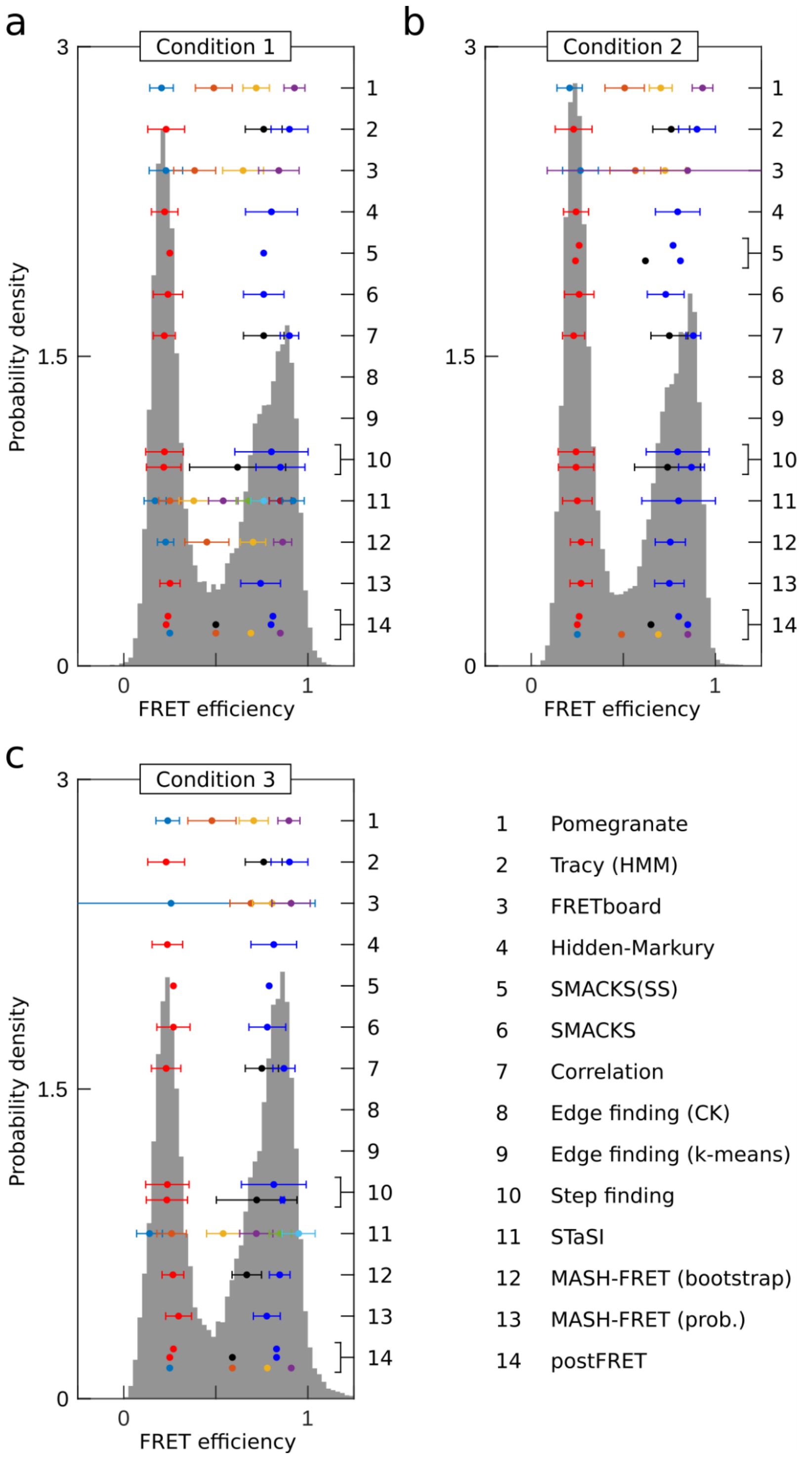


**Supplementary Figure 5 | FRET efficiency histograms and all inferred FRET states for the experimental datasets shown in Figure 5. a-c** The number of states and corresponding FRET efficiencies returned by the different analysis tools are shown for the experimental dataset shown in Fig. 5b,e,h, respectively. Brackets next to the right-hand tick markers indicate where multiple models with a different number of FRET states were submitted for a particular tool. The legend in c is valid for the entire figure (and throughout the paper).

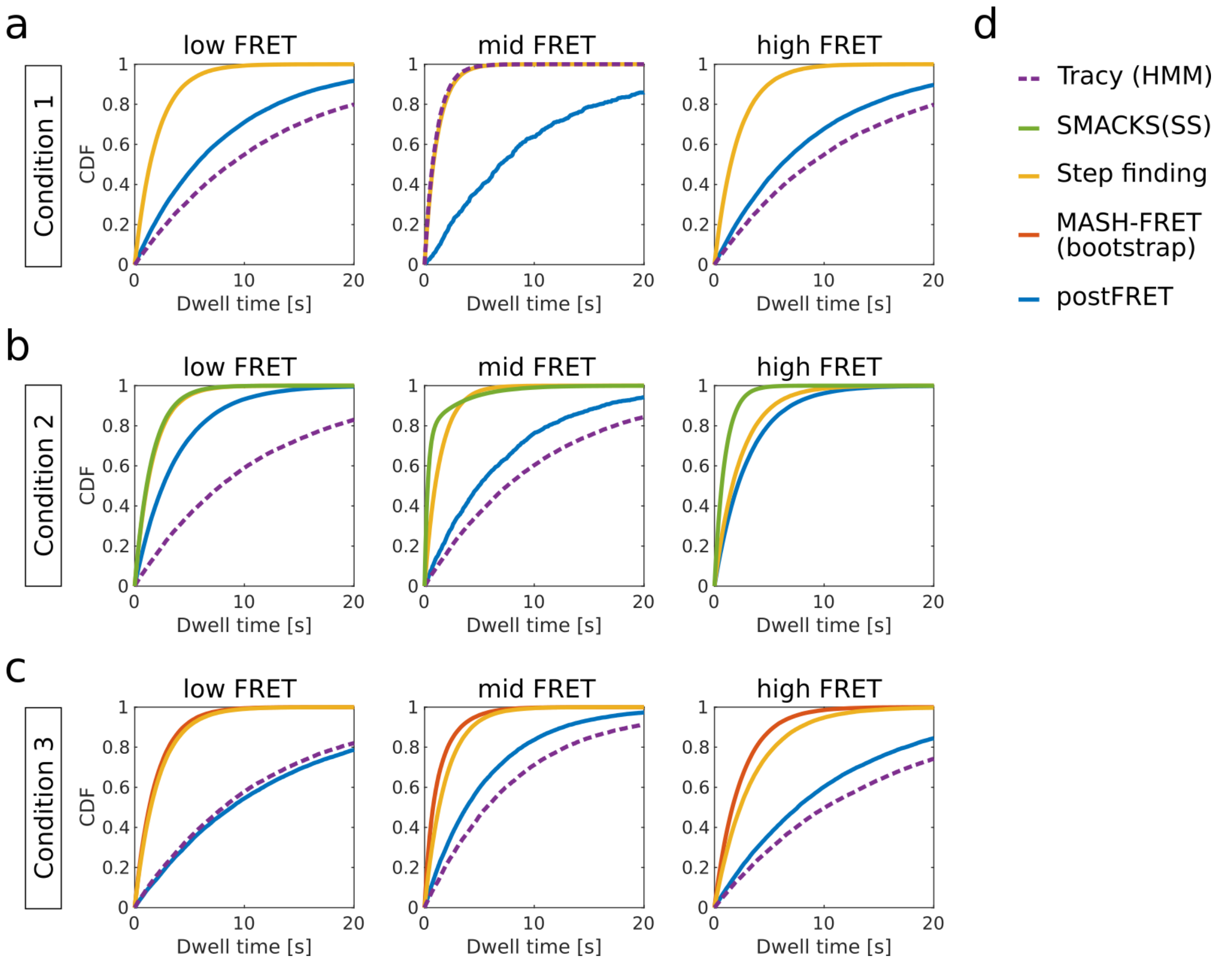


**Supplementary Figure 6 | Comparison of the kinetic models with three FRET states inferred for the dataset shown in Figure 5. a-c** Cumulative distribution function (CDF) of the dwell times simulated using the inferred kinetic models (described in Supplementary Note 3) are shown for conditions 1, 2, and 3, respectively. Please note the variation among the inferred FRET efficiencies of the low, mid and high FRET states (Supplementary Figure 5), hindering a direct kinetic comparison in this case. The model inferred by *Tracy* does not populate the high FRET state under condition 2, hence *Tracy* data is absent in the right panel in **b**.

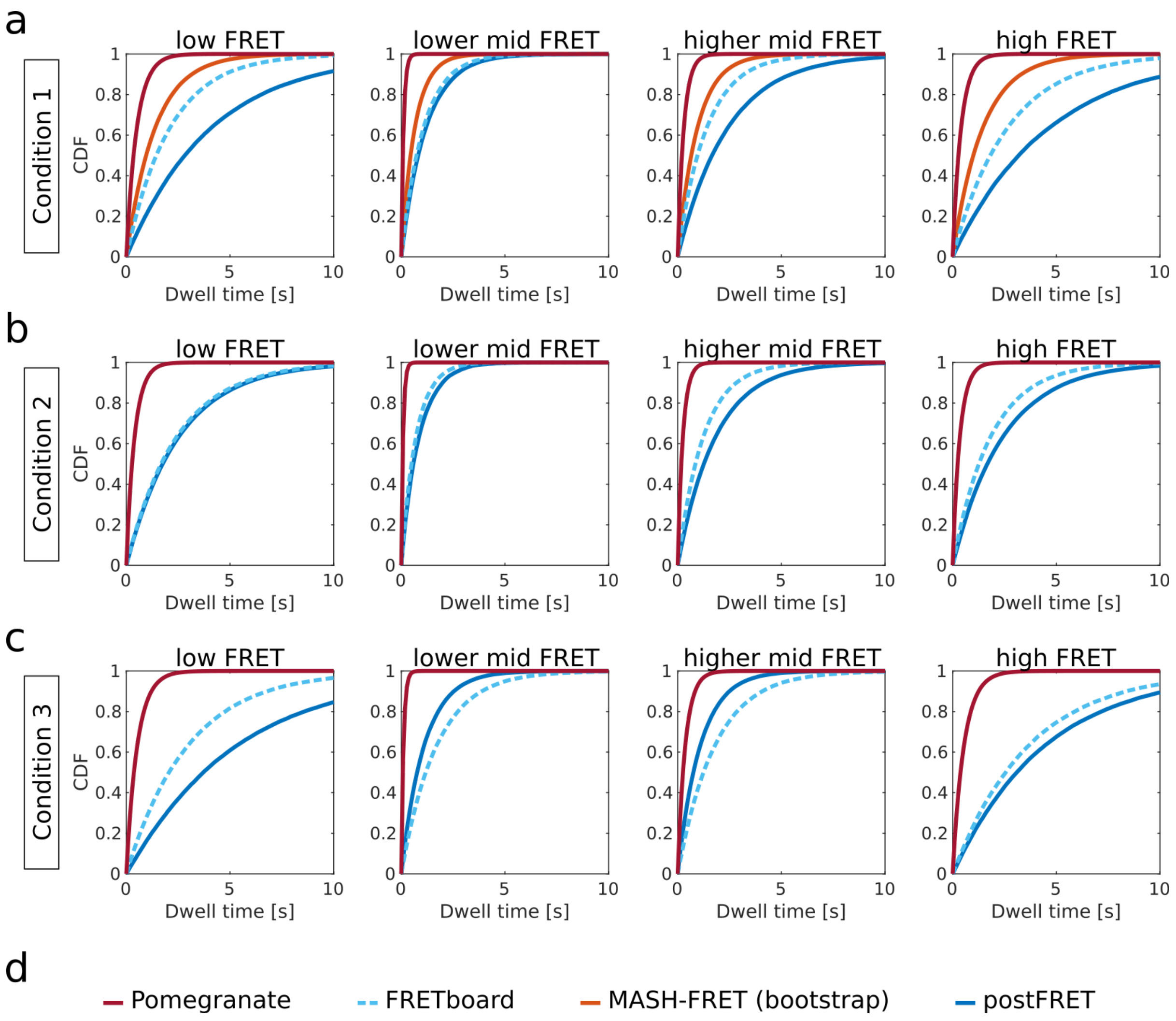


**Supplementary Figure 7 | Comparison of the kinetic models with four FRET states inferred for the dataset shown in Figure 5. a-c** Cumulative density function (CDF) of the dwell times simulated using the inferred kinetic models (described in Supplementary Note 3) are shown for conditions 1, 2, and 3, respectively. Please note the variation among the inferred FRET efficiencies of the four FRET states (Supplementary Figure 5), hindering a direct kinetic comparison in this case.

### 2 Supplementary Notes

**Supplementary Note 1: Simulation of smFRET trajectories**

In short, simulated smFRET datasets were generated to mimic fluorescence traces obtained by TIRF-based experiments. State trajectories were modeled with a continuous-time approach and later discretized. Similar to experiments, this allows state transitions to occur during the integration time window (time bin of the detector). Noise was added to the fluorescence intensity traces using experiment-derived parameters to generate realistic data.

In more detail, for each "molecule" a continuous-time state trajectory was simulated based on the kinetic model, as specified by a transition rate matrix. A summary of the specific simulation parameters is given in the Supplementary Table N1 and all configuration files with all parameters are provided as Supplementary Datafiles. First, the trace length was determined from an exponential distribution described by the rate of photobleaching. The trace length was rejected if it was shorter than a minimal trace length and truncated to a maximal trace length (see Supplementary Table N1). Then, a random initial state was chosen based on the probability of being in a particular state given the transition rate matrix. Starting from this state, dwell times for all possible transitions to the other states were drawn randomly from exponential distributions defined by the transition rates, and the shortest dwell time determined the transition and the new state of the system. This process was repeated until the full trace length was reached.

This state trajectory was then converted into discrete-time fluorescence intensity traces using a specified sampling rate. For each time bin (i.e., camera frame), the donor and acceptor intensities upon donor excitation and the intensity of the acceptor upon acceptor excitation were drawn from state-specific Gaussian distributions (specified by the means $\mu_I$ and covariance matrices given in the configuration file). The intensity in each channel during a time bin is given by the weighted average of all states visited during this specific time bin.

Typically, single-molecule fluorescence traces show variations in the fluorescence level between individual molecules, due to, amongst others, local variations in excitation power and local dye environment[1]. To take these variations into account, two additional sources of per-trace intensity variations were considered for the simulated data shown in Figs. 3 and 4. First, for each molecule, individual intensity levels for each state were chosen. To do so, the intensity level was drawn from an empirically determined state-specific Gaussian distribution (with mean $\mu_I$ and standard deviation $5 * \sqrt{\mu_I}$). Second, for each molecule, an individual brightness factor was determined by $1.20^r$ where $r$ was randomly chosen from the interval [-1, 1]. Thus, this factor is distributed in the interval [0.83, 1.20] and all channels were multiplied by the same factor. For the simulated data shown in Fig. 4, independent blinking of the donor and acceptor dye was modeled by a simple 2-state system ("bright", "dark"). In the case of an acceptor dark state, the FRET efficiency was set to zero. Details are given in Supplementary Table N1.

Five hundred additional datasets from the same parameter set were created and compared, to validate that the dwell time distribution of the dataset used in this study shows the expected behaviour (see Supplementary Fig. 3). Configuration files with all simulation parameters (including the ground truth for the kinetic models) for the synthetic data in Figs. 2, 3, and 4 can be found in the Supplementary Datafiles. The MATLAB scripts used for the simulation are publicly available at: [www.kinSoftChallenge.com](http://www.kinSoftChallenge.com) and [https://doi.org/10.5281/zenodo.5701310](https://doi.org/10.5281/zenodo.5701310) .

**Supplementary Table N1:** Parameters for the simulation of smFRET traces

| | **Fig. 2** | **Fig. 3** | **Fig. 4** |
|---|---|---|---|
| kinetic model (rate constants next to arrows are in $s^{-1}$) | 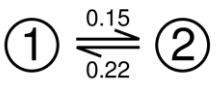 | 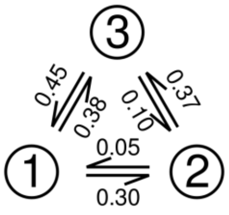 | 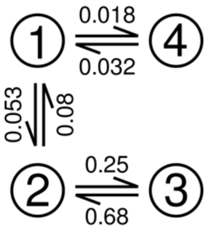 |
| sampling rate ($s^{-1}$) | 5 | 10 | 5 |
| bleach rate ($s^{-1}$) | 0.007 | 0.02 | 0.025 |
| min. trace length (s) | 50 | 10 | 8 |
| max. trace length (s) | 400 | 200 | 200 |
| kinetic heterogeneity? | no | no | yes |
| assignment state -> FRET level | ① -> low FRET<br>② -> high FRET | ① -> low FRET<br>② -> mid FRET<br>③ -> high FRET | ①, ② -> low FRET<br>③, ④ -> high FRET |
| blinking included? | no | no | yes<br>$k_{bright}$ = 7 $s^{-1}$<br>$k_{dark}$ = 0.007 $s^{-1}$ |
| per trace emission variability?[a] | no | yes | yes |
| per trace excitation variability?[a] | no | yes | yes |
| SNR (estimate)[b] | 4 | 3 | 4 |
| number of traces | 75 | 150 | 250 |

[a] The exact parameters can be found in the configuration files for the simulation in the Supplementary Datafiles.

[b] The SNR estimate is based on the separation and width of the peaks in the FRET efficiency histogram. Peaks were fitted with Gaussian distributions and the two peaks with minimal separation were considered. The SNR was then calculated by $|\mu_1 - \mu_2|/\sqrt{{\sigma_1}^2 + {\sigma_2}^2}$, where $\mu$ and $\sigma$ are the mean and standard deviation of the Gaussian functions, respectively. Exact parameters for the state-specific fluorescence intensities can be found in the configuration files for the simulation in the Supplementary Datafiles.

**Supplementary Note 2: Estimated minimal uncertainty of rate constants inferred from simulations**

Because of the finite number of traces per datasets, only a limited random sample of dwell times is observed for each given transition, resulting in a variation of the rate constants inferred from different datasets with identical ground truth. In order to estimate this lower bound of the uncertainty for the inference of rate constants from a finite dataset, we randomly drew the same number of dwell times as provided in the simulated challenge dataset from an exponential distribution with time constant $\tau = 1/k$. The maximum likelihood estimator (MLE) for the rate constant that produced this set of dwell times $\Delta t$ is given by $1/\overline{\Delta t}$. This calculation of the MLE was repeated one million times. The standard deviation of these 1 million MLEs is a function of the number of dwell times present in the challenge data set – the more dwell times are observed, the narrower the MLE distribution – and hence, it depends on the transition rate constants and the total observation time. We used this standard deviation as an estimate of the lower bound for the uncertainty of inferred rate constants from the simulated datasets.

**Supplementary Note 3: Simulation of cumulative dwell-time distributions from inferred kinetic models**

In order to compare submissions with the same number of FRET states but different underlying kinetic models (i.e., number of hidden states and connectivity), we simulated dwell times from the submitted kinetic models for the three datasets shown in Fig. 4 and 5. This yields cumulative dwell-time distributions that are characteristic for the kinetic model. Dwell times were accumulated from simulations of continuous time state trajectories (Supplementary Note 1) that included roughly 200x (Fig. 4d) or 400x (Fig. 5c,f,i) more time points than the original datasets.

**Supplementary Note 4: A simple file format for smFRET trajectories**

So far, no standardized and widely accepted file format for storing and exchanging smFRET trajectories is in use. A "single-molecule dataset" (SMD) file format has been proposed[2], based on JavaScript Object Notation (JSON), but has not been broadly adopted in the community. In this study, we opted for a simple tab-delimited text file format that is sufficient for the encountered time trace datafile sizes, circumvents intricate parsing, and was readily utilized by all participants of this study.

Each molecule is represented by a separate, tab-delimited text file. Each file contains a column with the time information and columns with the fluorescence intensities of the donor and acceptor after donor excitation ($I_{Dem|Dex}$, $I_{Aem|Dex}$). Additionally, columns with the acceptor intensity after acceptor excitation ($I_{Aem|Aex}$) and the apparent FRET efficiency $E_{app} = I_{Aem|Dex} \,/\, (I_{Aem|Dex} + I_{Dem|Dex})$ are present for the simulated dataset. The file format can be easily extended with additional columns for additional detection channels, e.g., more spectral and/or polarization channels. In addition to the time series data itself, further metadata, describing experimental conditions, acquisition parameters, and settings used to extract the intensities from the recorded raw data, can be included, either in the header for each tab-delimited file or in a separate file, as outlined in a recent position paper of the FRET community[3]. The broad acceptance attained in this study forms a promising starting point for the urgently needed dissemination of a common shareable file format for smFRET trajectories.

### 3 Supplementary Methods

All analysis tools are detailed here in the order of the numbering in the maintext.

##### Supplementary Method 1: Pomegranate

**Overview**

The workstream in pomegranate utilizes the fast and flexible probabilistic models built into the python package Pomegranate for efficient and iterative model formulation, fitting and evaluation using the Bayesian Information Criterion (BIC). The presented version of the workstream requires data preprocessing, where smFRET trajectories are sorted and only valid trajectories (based on expert valuation) are passed on through the analysis. Dwell time analysis is subsequently performed after defining all transitions using a Multivariate Gaussian fitting scheme and unbinned maximum likelihood fitting**.** All parts of the process can be evaluated and customized using user inputs based on expert evaluation and iteratively improved.
A solution to tedious manual data sorting has since this analysis been implemented in an “end-to-end” GUI software[4], that allows both sorting of smFRET trajectories based on deep learning and simultaneously applies the full analysis workstream presented and used here. The code used here as well as the compiled DeepFRET programme can be freely downloaded at www.hatzakislab.com/#software. The Deep Neural network sorting step is not necessary here as the data are presorted.

**Structure of analysis workstream**

Step 1: Model formulation
Accepted smFRET data is loaded into the Python workstream, where the FRET efficiency is calculated from donor and acceptor intensities and subsequently analyzed using Hidden Markov modelling. Initially the number of underlying FRET distributions should be determined to optimize the model fit. To extract this information, the software allows the user to fit all data to a Hidden Markov model containing between 1-n gaussian distributed state populations (shown to be a robust approximation of FRET distributions[5]) and corresponding transitions between them using and the Baum-Welch forward-backward algorithm. The models can then be compared and the best selected for further analysis using Bayesian Information Criterion (BIC) to penalize overfitting as shown earlier[6,7]. Once a model is accepted and finalized, it can be saved for further use and evaluation.

Step 2: Trace by trace prediction
A finalized model can be readily applied to analyze smFRET trajectories. The software will for each individual trajectory use the Viterbi algorithm to calculate the most likely state for each observation based on the provided model. To further validate model predictions, a subset of trajectories can be visualized with the corresponding idealized state (model prediction) for user evaluation and validation. Based on expert knowledge, users may choose to reiterate Steps 1 and 2 for optimal fitting of smFRET data. For each data point, the most likely state is found and all data is saved to allow any further analysis or visualizations.

Step 3: Parameter extraction and evaluation
Using the model predictions, each transition in each trajectory (E -> E+1) is plotted in a Transition Density Plot (TDP) and separated using a Gaussian Mixture Model (GMM) fitting scheme. The number of clusters was determined using a combination of user inspection of unfitted data and BIC evaluation. For each fitted cluster, only data points within a 99 % confidence interval of the cluster center would be included for kinetic rate extraction. This ensures tighter clusters and fewer single “off-cluster” points.

To extract kinetic transition rates, the dwell times of each cluster were fitted using a single exponential decay and maximum likelihood fitting, subsequent comparison to a two-component exponential decay using BIC was used to check for degenerative states.

**Supplementary Method 2: Tracy**

The *Trace Intensity Analysis* toolbox, referred to as TRACY, was programmed in MATLAB and has been updated to run on MATLAB R2018a. The program is used for analyzing two-color single molecule FRET Traces. The program, written by Gregor Heiss[8], performs the following tasks:

1. Extraction of single-molecule intensity trajectories for one or two channels with only donor excitation or with Alternating Laser Excitation from image stacks.
2. Allows manual selection of the frames providing useful fluorescence intensity data for each molecule.
3. Allows correction of the fluorescence intensity for determination of FRET values.
4. Allows user-defined categorization of smFRET traces and analysis of molecular subpopulations.
5. Analysis features includes a Hidden Markov Model analysis using the HMM toolbox in MATLAB written by Kevin Murphy (https://www.cs.ubc.ca/~murphyk/Software/HMM/hmm.html). The HMM analysis can be run individually on each trace or globally on an entire dataset.
6. From the HMM analysis, a Viterbi path can be calculated for each molecule and a transition density plot calculated.

Workflow:
For the analysis performed in this study, the following workflow was followed.
1. Data were loaded into TRACY and categorized manually by framewise selection of valid smFRET regions. Unselected regions in each trace were treated as bleached frames and were not used for further analysis.
2. Next, a global HMM analysis was performed on each data set. For the HMM analyses, only the FRET efficiency data were used and not the donor and acceptor intensities. The mean FRET efficiency and sigma were set as learning parameters.
3. A Viterbi path was then calculated from the given HMM parameters for the individual traces. Using the determined transitions, a transition density plot (TDP) between the learned states was determined.
4. From the TDP, individual states were manually selected and the corresponding dwell-time histograms were fitted with single- or double-exponentials to obtain the transition rates.

In the end, the decision regarding which model to apply and analysis of the returned dwell-time distributions were determined manually based on visual inspection of the data and results from the initial analysis. The time involved varied from 20 to 60 min depending on the size and complexity of the data set.
TRACY is available upon request.

#### Supplementary Method 3: FRETboard

FRETboard is a semi-supervised FRET trace classification tool that is served remotely through a web browser. Using a simple click-and-point interface, the user may 'teach' a classification model to recognize certain patterns in the traces, by iteratively performing manual curation on an automatically classified example trace and then retraining the model using the corrected traces. This lends FRETboard the flexibility to easily adapt to different labeling schemes. To further expand this flexibility, the user may also experiment with different combinations of nine features derived from the original channels, and the application of different model structures. These properties and further details on FRETboard usage as applied in this challenge are described below.

##### Model structures

While essentially any supervise-able classification model type may be trained through FRETboard, three flavors of hidden Markov models (HMMs) are currently included by default. The "vanilla" structure produces a straightforward fully connected HMM sporting no further modifications. The "boundary-aware" structure adds additional "edge states" between states, which are trained on measurements around a detected transition. Transitions between states may only occur through these edge states. If state transitions are marked by a signature distribution in a certain feature, this distribution is captured by the edge state, which allows for more accurate detection of state transitions. The "GMM-HMM" structure also implements edge states, and in addition models emissions using a Gaussian mixture model (GMM), which adds the flexibility to classify noisier distributions as a single state using multiple Gaussians. The number of Gaussians per GMM is determined per state using a Bayesian information criterion selection procedure. Users may also write a custom model structure implementation, using the provided template.

In this challenge, the vanilla structure was used for the two-state simulated data, while the GMM-HMM structure was used for the other data sets, to account for added noise and other complications.

##### Features

As different patterns may be better discernable using different features, users may activate and de-activate each of the nine included features as they see fit. These features include the original acceptor and donor channels, and the acceptor and donor channels during direct acceptor excitation if alternating laser excitation (ALEX) is employed. The proximity ratio $E_{PR}$ is included as an approximation of FRET efficiency and is defined as:

$$E_{PR} = \frac{F_{Aem}^{Dex}}{F_{Aem}^{Dex} + F_{Dem}^{Dex}}$$

Here, $F_{Aem}^{Dex}$and $F_{Dem}^{Dex}$are the original donor and acceptor emission. The summed intensity $F_{sum}$ is also included:

$$F_{sum} = F_{Aem}^{Dex} + F_{Dem}^{Dex}$$

Furthermore, we used two time-aggregated features that capture the variability of features over a sliding window of five measurements: the Pearson correlation coefficient between $F_{Aem}^{Dex}$ and $F_{Dem}^{Dex}$and standard deviation of $F_{sum}$. These features may aid models in capturing feature distributions characteristic for state transitions. For this challenge, training was started using the default combination of features ($E_{PR}$, $I_{sum}$, standard deviation of $I_{sum}$ and correlation coefficient). However if training accuracy failed to reach levels above 95%, the best functioning combination of features was picked, by gauging the effect of toggling features on training accuracy.

##### Training procedure

The procedure is initialized by fitting an unsupervised model on all loaded traces in a traditional manner, using randomly generated initial parameters that are then fitted using an implementation of expectation maximization. The only user-provided guidance at this point is the number of states that should be recognized. Traces are classified, and the trace classified with least certainty, i.e. for which the state path probability normalized over sequence length was lowest, is presented to the user for manual correction of the classification. The probability of assigned states for a given trace may be poor in this trace because of the presence of noise. If this is the case, a user may choose to assign noisy measurements to a state they deem appropriate. However, if it is more appropriate to remove the noise, as is the case in bleaching and blinking events, the user may filter these measurements out by assigning them to a separate state reserved for such events. Such a state will then be discarded before FRET distribution and transition rate analysis. Alternatively, the model fit may suffer if the trace contains more or fewer states than those included in the current HMM. In that case, the number of states and classification must be adjusted appropriately.

After applying manual corrections, the first semi-supervised training round on all loaded traces is started. State distributions and transition rates can now be deduced from the corrected trace and be used as initial parameters, after which the HMM is refitted on supervised and unsupervised traces simultaneously using semi-supervised expectation maximization. After refitting, traces are reclassified, and the trace now marked by the lowest state path probability is presented to the user. The procedure is repeated until the user finds that presented traces are correctly classified.

For this challenge, we chose to manually classify five traces and retrain the model after each manual classification. Analysis typically took 15 minutes per data set and was performed using the remote server version running at www.bioinformatics.nl/FRETboard.

**Parameter extraction**

FRET distributions are extracted by separating values from classified traces by state and calculating mean and standard deviation for each state. To obtain transition rates (*F*), a transition matrix (*A*) is derived from the classified data, which must then be converted from discrete to continuous rates and corrected for framerate ($f_s$):

$$F = I + f_s \times log\, A$$

Here $I$ is the identity matrix and $log$ denotes the natural matrix logarithm operation. 95% confidence intervals of transition rates are estimated by bootstrapping the classified traces and calculating *F* 100 times.

#### Supplementary Method 4: Hidden-Markury

Hidden-Markury is a trace analysis software based on a global optimization of one global kinetic model. It supports the global analysis of 1D FRET efficiency traces and 2D donor & acceptor photon streams with multiple model optimization options, such as degenerate states, forbidden transitions, and fixed model parameters. The core of the Hidden-Markov model and its optimization is based on the open python library hmmlearn (https://hmmlearn.readthedocs.io/en/latest/index.html)

The code of this software tool and its description is provided as interactive Jupyter notebooks and can be found in the GitHub repository (https://github.com/ChristianGebhardt/Hidden-Markury). Hereby, the notebook combines the high flexibility of individual code adaption and the user-friendly nature of interactive GUI elements with informative descriptions.

##### Step 0: Installation and Getting Started

The usage of the Hidden-Markury software requires the cloning of the GitHub repository (https://github.com/ChristianGebhardt/Hidden-Markury) and the installation of the python packages (hmmlearn, numpy, pandas, scipy, matplotlib) as described in the README file in more detail. The repository provides exemplary data sets to get started with.

##### Step 1: Data Import

The data can be imported from various delimited data formats (.csv, .tsv, etc.) where the user only needs to specify the columns/rows of the different data sources from donor (DD) / acceptor channel (DA) or the FRET efficiency (E). The time information is automatically extracted from the first entry.
The imported traces are visualized for manual inspection.

##### Step 2: Model Specification and Initialization

The Hidden-Markury notebook provides the options for 2D-trace analysis of donor (DD) and acceptor (DA) photon streams or 1D-trace analysis of the FRET efficiency traces. In both selected cases the states are initialised by a multi-Gaussian fit in the 2D/1D histogram, where the user needs to select the number of (non-degenerated) states (see Supplementary Method Figure 4.1).

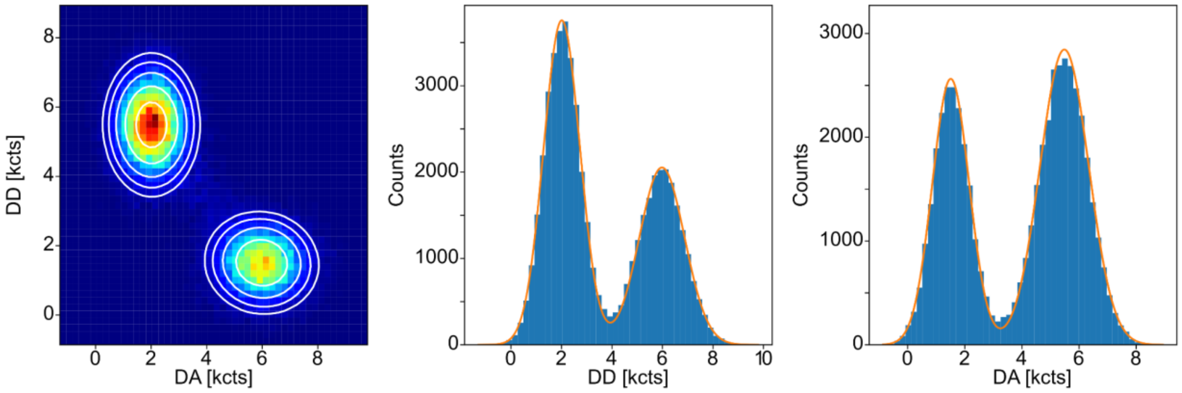


**Supplementary Method Figure 4.1:** 2D-histogram of donor (DD) and acceptor photon counts (DA) for all time traces fitted with two 2D-Gaussian distributions (left). 1D-projections of the 2D fit results and the 1D-histograms of DD (center) and DA (right).

**Step 3: Model Fitting and Prediction**

The notebook allows to manually adapt the model fitting procedure by fixing values such as the Gaussian emission functions, the transition matrix or individual forbidden transitions during the model optimization. For degenerated states, the initial fit values from step 2 are required to be fixed in the model fitting step. The actual model optimization uses the expectation-maximization (EM) algorithm[9] for a global optimization of all traces. In this study all values (Gaussian emission distributions, transition matrix, initial distribution) were optimized during this step. For the degenerated states, only the transition matrix and initial distribution were optimized.

The Viterbi-algorithm is used for the prediction of the states in all traces based on the optimized model (see Supplementary Method Figure 4.2).

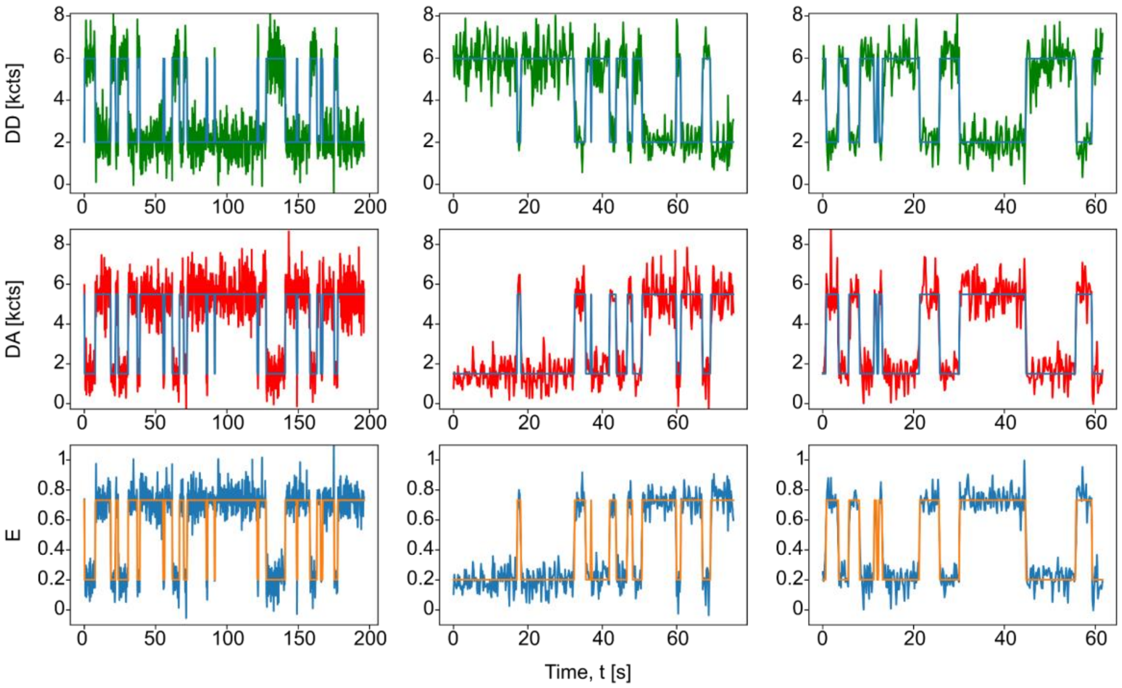


**Supplementary Method Figure 4.2:** Three exemplary traces with the donor (DD) and acceptor photon counts (DA) in the top and center row, respectively, and the calculated FRET efficiency trace in the bottom row together with the predicted photon counts (blue) and FRET efficiencies (orange).

**Model Assessment**

The optimal model can be assessed by multiple runs of step 2 and 3 with varying numbers of states (and potential degeneration) and assess the log likelihood of the prediction.

For the error estimation of the rate constants, the model optimization is repeated with a subset of all traces selected by sampling with replacement. The final results (mean and standard deviation) of the transition rate constants reported in this study are taken from a sampling of 40% of the traces and 20 repetitions.

**Supplementary Method 5: SMACKS**

Single molecule time traces hold valuable information about a protein's thermodynamics and kinetics. However, their analysis is far from trivial – especially, when their interpretation goes beyond apparent observations. *Single Molecule Analysis of Complex Kinetic Sequences* (SMACKS) uses mathematical models for pattern recognition to not only resolve statistically relevant rates from such traces but also their uncertainties[1]. Thereby, SMACKS does not rely on dwell-times but takes every experimental data point into account to optimise one global kinetic model. Consequently, it allows for experimental variation between individual trajectories[1]. At the core of SMACKS' analysis lie hidden Markov Models that establish a mathematical relation between experimental observations and their subsequent interpretation, which in turn is limited by a predefined number of states[10] . The general procedure of how SMACKS was used for data analysis within this study will be described in the following. A more detailed description on how to use SMACKS can be found in the associated manual (https://www.singlemolecule.uni-freiburg.de/software/smacks).

**Step 1: Software Installation and Data Import**
Being implemented in Igor Pro (Wavemetrics), SMACKS' source code was downloaded from https://www.singlemolecule.uni-freiburg.de/software/smacks. The program was started using the *startSMACKS* shortcut. To load the kinSoftChallenge data files, the dataID in the startSMACKS.ipf script editor was changed to StrConstant dataID = "time;g_g;r_g;r_r;fret;". They were imported by using the ascii importer in the SMACKS menu (**-->Import ascii**). Afterwards, as SMACKS only accepts files with up to three tab- or space-separated data columns, the dataID was changed back to StrConstant dataID = "g_g;r_g;r_r" for further analysis.

**Step 2: Trace-by-Trace HMM (TbT)**
The TbT workflow was started in the SMACKS menu (**-->Init TbT**). In the settings, the number of apparent states was assigned by eye according to the user's observation. Adjustments as such were confirmed by clicking **Initialize**. To assign each trace a Viterbi path (meaning certain states according to the Viterbi algorithm), the procedure was applied to the whole dataset by choosing **-->TbT Batch Converge** in the SMACKS menu. Next, all associated Viterbi paths were individually checked and approved by browsing through the traces (using **<<** and **>>**) and deleting parameters causing inappropriate Viterbi paths (using **Del**). For traces that were assigned a Viterbi path despite not reaching all states, the parameters were deleted as well. Eventually, by using **TbT Apply Means**, the mean of all saved (i.e. correct) parameters was applied to the remaining traces.

**Step 3: Semi-Ensemble HMM (ENS)**
Afterwards, the ENS workflow was started in the SMACKS menu (**-->Init ENS**). Here, different state models were analysed based on the apparent states and optimised parameters found in the TbT workflow. Depending on the possibility of hidden states (i.e. states displaying the same FRET efficiencies while differing kinetically), different state configurations were tested. Different states were indicated by different numbers (0, 1, 2, etc.), whereas associated hidden states were denoted by doubling them (e.g., 001, 011, 0011, etc). Using the add-on feature **-->Compare BICs**, the state model that best represented the data (i.e. the one with the lowest BIC) was chosen. Confidence intervals for the chosen state model were calculated using the feature **-->Confidence Intervals** in the SMACKS menu.

**Step 4: Calculating Kinetic Rates and FRET Efficiencies**
From the **-->TbT Apply Means** procedure after the TbT workflow, the mean intensity values over all donor (D) and acceptor (A) traces were obtained. They were used to calculate the FRET efficiencies E according to:

$$FRET\ E = \frac{A}{A+D} \tag{5.1}$$

Corresponding uncertainties were calculated according to:

$$\sigma = \sqrt{\left(\frac{D}{(A+D)^2}\right)^2 \cdot \sigma_A^2 + \left(\frac{-A}{(A+D)^2}\right)^2 \cdot \sigma_D^2} \tag{5.2}$$

In the ENS workflow, SMACKS calculated a transition probability matrix and a covariance matrix of the Gaussian probability densities for all states. This calculation is performed in user-supplied time units. Therefore, giving specific time information is not necessary: the transition probabilities $a_{ij}$ from state i to j are specified for this given time interval. Converting those probabilities from the transition probability matrix to rate constants $k_{ij}$ (Hz) was done by multiplying them with with the sampling rate (frames/second) according to:

$$kinetic\ rate\ k_{ij} = transition\ probability\ a_{ij} \cdot sampling\ rate \tag{5.3}$$

Confidence Intervals were converted to Hz accordingly.

**Parameter Settings, Technical Specifications and Computation Time:**
In steps 2 and 3, specific parameters can be set and varied. The initial parameters in the TbT workflow, however, were not adjusted. Instead, the given initial values were used. To find out which state model was most likely, the default number of iterations (500) given in the ENS workflow was halved to save computation time. Then, for a more precise calculation of the transition probabilities of the chosen model, the default values for the number of *Max. Iterations* (500) as well as for the *Convergence Threshold* (1E-15) were kept. However, this affected the values in the transition matrices only within the margins of error.

The synthetic datasets were analysed on a MiFcom desktop computer (8.00 GB RAM, Intel(R) Core(TM) i5-4460 CPU@3.20 GHz, 64-bit operating system, Windows 10). The experimental sets were analysed on a MiF desktop computer (32 GB RAM, Intel(R) Xeon(R) CPU E5-2650 v3 @ 2.30 GHz 2.30 GHz (2 prozessors), 64-bit operating system, Windows 10). In both cases, Igor Pro 6.37 was used.

The computation time varied depending on the complexity and size of the analysed set and the computer used for the analysis. The first synthetic data set containing a two-state system without hidden states was analysed within 30 minutes. The other two sets, in contrast, contained hidden states and – in case of the third set – more than two apparent states. Therefore, the second set required 4 hours, whereas the third took a little less than 13 hours to be analysed completely.

For the experimental sets, computation time depended on time resolution. Under 10 kHz conditions, data analysis took 35 hours, whereas the 0.1 kHz data set was analysed within 1.5 hours.

**Supplementary Method 7: Correlation**

**Introduction**

We recently presented a quantitative model for fluorescence correlation curves of complex multi-state kinetic networks obtained in single-molecule FRET experiments using MFD[11]. Here, we extended this methodology for use with fluorescence traces of immobilized molecules. In principle, this simplifies the analysis by removing the diffusion term of the correlation function[12–14], but modifications are required to obtain accurate correlation functions from variable length traces. In MFD experiments, we are further able to compute filtered correlation functions by utilizing e.g. the lifetime information[14–16], but no such information is available in the given case. To this end, we applied a step-finding algorithm to convert the observed FRET efficiency traces into digitized state trajectories, which were further used to compute filtered correlation functions. In the ideal case, this allows us to resolve the kinetics even in multi-state networks of three or more interconverting states. While an accurate estimation of the microscopic rate constants for the more complicated cases involving degenerate FRET states was not possible, correlation analysis still offers a minimally biased approach to estimate the kinetic relaxation times of the corresponding transition rate matrix, which allows to validate kinetic models inferred by other methodologies.

**Methods**

All analysis was done in MATLAB. The code and all analysis files are available at https://doi.org/10.5281/zenodo.5512005.

**Computation of correlation functions**

The calculation of unbiased correlation functions from the variable-length fluorescence time traces required three problems to be addressed:

1. Correlation functions must be calculated for the total duration of the traces as dynamics and trace length may occur on a similar timescale. Edge-effects to the lower sampling of long lag times thus need to be accounted for.
2. Correlation functions obtained from traces of different lengths must be correctly averaged.
3. Errors arising due to trace-by-trace variability must be estimated.

The correlation function between two signals $S_A(t)$ and $S_B(t)$ is defined by:

$$G_{AB}(\tau) = \frac{\langle S_A(t) S_B(t+\tau)\rangle}{\langle S_A(t)\rangle\langle S_B(t+\tau)\rangle}, (7.1)$$

where $\tau$ is the lag time and $\langle \dots \rangle$ denotes the time average for a long measurement. Generally, the correlation function is calculated only up to a time lag that is a fraction of the total measurement time ($\tau \ll T$, typically up to a maximum of 1/10 of the measurement time). To compute the correlation function over the whole length of the trace, it was calculated for every trace $k$ as described in references [17,18] by:

$$G_{k,AB}(\tau) = \frac{(T_k - \tau)\sum_{t=0}^{T_k-\tau} S_A(t) S_B(t+\tau)}{\sum_{t=0}^{T_k-\tau} S_A(t) \sum_{t=\tau}^{T_k} S_B(t+\tau)}, (7.2)$$

where $T_k$ is the length of the trace and the sums only extend over the valid ranges of the time trace, i.e. $0 \leq t \leq T_k - \tau$ for $S_A(t)$ and $\tau \leq t \leq T_k$ for $S_B(t)$.

Due to the variable length of the traces, the average correlation function over multiple traces is not simply equivalent to the average of all trace-wise correlation functions. To compute the average correlation function over all traces, the terms in the above equation were computed for every trace $k$ and the final correlation function over all traces was computed as:

$$G_{AB}(\tau) = \frac{\sum_k (T_k - \tau)\left[\sum_k \sum_{t=0}^{T_k-\tau} S_A(t) S_B(t+\tau)\right]}{\left[\sum_k \sum_{t=0}^{T_k-\tau} S_A(t)\right]\left[\sum_k \sum_{t=\tau}^{T} S_B(t+\tau)\right]}. (7.3)$$

Correlation functions were calculated for lag times on a multiple-tau scheme over stretches of linear time lags with exponentially increasing spacing, i.e. $\tau = 0,1,2,\dots,19,20,22,24,\dots,38,40,44,48 \dots$ etc (see reference[19] for details).

To estimate the error due to trace-by-trace variability, we performed bootstrapping. For a set of $N$ traces, we randomly selected $N$ traces with replacement (that is, duplicate selections are allowed) 50 times, computed the average correlation function $G_{AB}$ according to eq. 7.3, and determined the standard error of the mean from the set of correlation functions.

FRET-FCS correlation functions were calculated from the background-corrected intensities in the donor and FRET channels to compute the auto- and cross-correlation functions of the donor (D) and acceptor (A) signals ($G_{DD}$, $G_{AA}$, $G_{DA}$ and $G_{AD}$). Filtered-FCS curves were computed from the digitized state trajectories determined from the step-finding analysis (see below). For the analysis, all possible auto- and cross-correlation functions between the identified states were used (i.e., 4 in the case of two FRET states and 9 in the case of three FRET states).

**Estimation of FRET efficiencies of the different states**

The minimal number of states and their FRET efficiencies are determined by Gaussian fitting of frame-wise FRET efficiency histograms. We used a Gaussian mixture model as implemented in the MATLAB function *fitgmdist*, based on an iterative Expectation-Maximization algorithm of the likelihood function. An exemplary fit is shown in Supplementary Method Figure 7.1A.

**Step-finding algorithm**

We apply the algorithm developed by Aggarwal et al. to identify steps in the FRET efficiency traces[20]. The algorithm does not assume any particular kinetic scheme but estimates the optimal number of steps based on the noise of the signal. Overfitting is avoided by introducing a penalty for each transition. For the analysis, we set an estimated noise of based on the distribution width obtained from the Gaussian fitting analysis ($\sigma_E$ = 0.05-0.1). No restraints are imposed on the FRET efficiencies of the steps. An exemplary result of the step-finding is shown in Supplementary Method Figure 7.1C. After the step finding, the stepwise FRET efficiency histograms were manually examined to identify thresholds to digitize the FRET efficiency trajectory (see Supplementary Method Figure 7.1B).

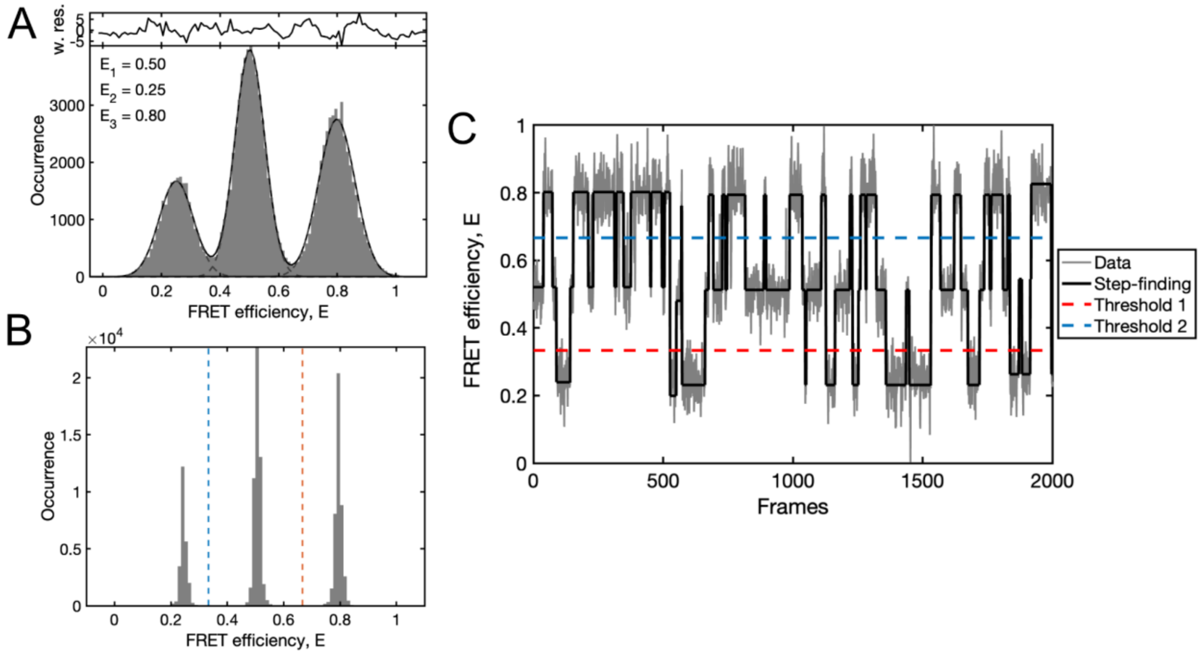


**Supplementary Method Figure 7.1: Exemplary workflow for the correlation analysis of single-molecule time traces. A-B)** FRET efficiency histograms of test dataset 2 before (A) and after (B) applying the step finding. Dashed lines in B indicate the used thresholds to define the state trajectory. **C)** Example of the step-finding algorithm. The idealized signal trajectory (black) is estimated from the noisy data (gray). Thresholds for the digitization of the FRET efficiency trajectory into states are shown as dashed lines.

**FCS model functions**

Depending on the complexity of the datasets, three different analyses approaches were used throughout this study, depending on whether the FCS curves were calculated from the fluorescence intensities (FRET-FCS) or the digitized state trajectories (filtered-FCS). Further, for cases where insufficient information could be inferred to decide on the number of kinetic states *a priori*, an empirical model was applied to estimate the kinetic relaxation times without consideration of the correlation amplitudes.

***FRET-FCS***

For two-state dynamics, analytical functions are known for the color-FCS curves:[4]

$$G_{DD}(\tau) = \frac{k_{12}k_{21}\Delta E^2}{(k_{12}(1-E_1)+k_{21}(1-E_2))^2}e^{-(k_{12}+k_{21})\tau} \quad (7.4)$$

$$G_{AA}(\tau) = \frac{k_{12}k_{21}\Delta E^2}{(k_{12}E_1+k_{21}E_2)^2}e^{-(k_{12}+k_{21})\tau} \quad (7.5)$$

$$G_{DA}(\tau) = G_{AD}(\tau) = -\frac{k_{12}k_{21}\Delta E^2}{\big(k_{12}(1-E_1)+k_{21}(1-E_2)\big)(k_{12}E_1+k_{21}E_2)}e^{-(k_{12}+k_{21})\tau} \quad (7.6)$$

Here, $E_1$ and $E_2$ are the FRET efficiencies of the two states and $k_{12}$ and $k_{21}$ are the rates of going from state 1 to 2 and backwards, respectively. For the FRET-FCS analysis, the FRET efficiencies of the states were fixed to the values determined by the analysis of the FRET efficiency histograms.

***Filtered-FCS***

In filtered-FCS, the correlation functions represent the pure correlation functions between the different states. For the two-state case, the analytical correlation functions between states 1 and 2 are then given by:

$$G_{11}(\tau) = \frac{k_{12}}{k_{21}}e^{-(k_{12}+k_{21})\tau} + c \quad (7.7)$$

$$G_{22}(\tau) = \frac{k_{21}}{k_{12}}e^{-(k_{12}+k_{21})\tau} + c \quad (7.8)$$

$$G_{12}(\tau) = G_{21}(\tau) = -e^{-(k_{12}+k_{21})\tau} + c, \quad (7.9)$$

where $c$ is a constant offset. The correlation functions are calculated from the matrix exponential of the transition rate matrix, $e^{K\tau}$, which can be obtained from the eigen-value decomposition of $K$:

$$K = \sum_{i=0}^{n-1} \Gamma_i\lambda_i \quad \Rightarrow \quad e^{K\tau} = \Gamma_0 + \sum_{i=1}^{n-1} \Gamma_i e^{\lambda_i t_c} \quad (7.10)$$

where $\lambda_i$ are the eigen-values and $\Gamma_i$ the eigen-matrices of $K$. A transition rate matrix of dimension $n$ has $n-1$ non-zero eigen-values, and the zero-th eigenvalue, $\lambda_0 = 0$, can be neglected in this case. The full correlation function is then obtained as:

$$G_{ab}(\tau) = \frac{\sum_{i=1}^{n-1} S_a^T \Gamma_i x_d S_b e^{\lambda_i\tau}}{\langle S_a\rangle\langle S_b\rangle} \quad (7.11)$$

where $S_a$ and $S_b$ are the state vectors, i.e. $\left\{\begin{pmatrix}1\\0\\0\end{pmatrix},\begin{pmatrix}0\\1\\0\end{pmatrix},\begin{pmatrix}0\\0\\1\end{pmatrix}\right\}$, $x_d$ is the vector of equilibrium fractions of the states and $\langle S_a\rangle$ and $\langle S_b\rangle$ are the average occupancies of the states, $\langle S_a\rangle = x_a$. Using this formalism, we can directly fit all rate constants of the transition rate matrix to the obtained correlation functions. A detailed derivation of the correlation functions is given in reference[11]. The filtered-FCS curves were analyzed globally with respect to the transition rate matrix.

***Empirical model functions for degenerate cases***

For the advanced cases where degeneracy of FRET states is involved (i.e., states with identical FRET efficiencies but different kinetic properties), our method cannot resolve the kinetics accurately. However, it is still possible to determine the relaxation times of the kinetic process from the correlation functions using simplified multi-exponential model functions with $n$ components.

$$G_{11}(\tau) = \sum_{i=1}^{n} A_{11,i}e^{-\frac{\tau}{\tau_i}} + c \quad (7.12)$$

$$G_{22}(\tau) = \sum_{i=1}^{n} A_{22,i}e^{-\frac{\tau}{\tau_i}} + c \quad (7.13)$$

$$G_{12}(\tau) = G_{21}(\tau) = -\sum_{i=1}^{n} A_{12,i} e^{-\frac{\tau}{\tau_i}} + c\,, \quad \text{where} \sum_{i=1}^{n} A_{12,i} = 1. \quad (7.14)$$

Here, $c$ is a constant offset. Here, to reduce the number of free fit parameters, we took advantage of the fact that the cross-correlation functions contain the same information and that the amplitudes should sum to 1 for the filtered-FCS curves.

The obtained relaxation times $\tau_i$ correspond to the inverse of the negated eigenvalues of the transition rate matrix, $\lambda = eig(K)$, and can thus be compared to the input values:

$$\tau_i = -\lambda_i^{-1}. \quad (7.15)$$

No further interpretation of the obtained amplitudes is performed in this case. All relaxation times were globally optimized over all correlation functions.

**Curve fitting**

Curve fitting was performed based on the reduced chi-square $\chi_r^2$ defined as:

$$\chi_r^2 = \frac{1}{N-k} \sum_i \frac{\left(G_{data,i} - G_{model,i}\right)^2}{\sigma_i^2}, \quad (7.16)$$

where $N$ is the number of data points, $k$ is the number of parameters of the model and $\sigma_i$ is the estimated error. Optimization was performed using the *fminsearch* function of MATLAB using the Nelder-Mead method[21]. For model selection, we applied the Bayesian information criterion[22] given by:

$$BIC = k\, ln(N) + \chi^2, \quad (7.17)$$

whereby the model with the lowest value for the BIC was chosen. Errors of the fitted parameters were estimated based on Metropolis-Hastings sampling of the posterior probability distribution of the model parameters using a flat prior[23,24]. The Markov chain Monte Carlo sampler as implemented in the *mhsample* function of MATLAB was run for 10.000 iterations using a symmetric proposal distribution. The width of the proposal distribution was estimated based on the errors of the fit parameters determined from the non-linear least squares fitting using the Hessian matrix at the solution, and set to one-tenth of this value. Every hundredth sample of the Markov chain was kept and used to calculate the confidence intervals. All given errors are specified as 95% confidence intervals. An example of the posterior distribution obtained for the elements of the transition rate matrix of the synthetic three-state system is shown in Supplementary Method Figure 7.2B.

**Performance considerations**

On a standard desktop computer, the calculation of FRET-FCS correlation curves for the datasets used in this study took less than a minute. For the filtered-FCS analysis, the limiting step is the step-finding algorithm, resulting in typical computation times of 10-15 minutes. Curve fitting generally took less than one minute. However, depending on how many models were tested, the total analysis procedure including human intervention could take up to 1 h.

**Analysis of the different datasets**

Correlation functions for all synthetic datasets were obtained by the filtered-FCS workflow. The correlation functions for the synthetic datasets of level 1 and 2 were analyzed using the analytical model functions for a two- and three-state system, respectively. Due to the degeneracy of the FRET states for level 3, the empirical analysis was performed with a three-component model. The number of components was hereby estimated by comparing the BIC of the models with two to four components.

Due to the high number of FRET states in the first experimental dataset, the empirical analysis with a two-component model was performed. For the second experimental dataset, the filtered-FCS workflow was applied with a two-state kinetic model for the 10-ms dataset. In addition, the correlation functions revealed a slower process on the minute timescale. Two estimate the timescales of these dynamics, the empirical analysis with a two-component model was additionally applied to the filtered-FCS curves. For the 1-ms dataset, only FRET-FCS analysis could be performed as the step-finding algorithm could not be applied to the noisy traces.

**Deviations of the correlation approach for the three-state system (Synthetic Data, Level 2)**

The large deviations of the correlation approach for the synthetic three-state system were a partly surprising result as the method had worked better in previous benchmarks. Based on the ground-truth transition rate matrix of the three-state system (see Supplementary Datafiles), the two relaxation times of the simulated system are $\tau_1$ = 1.07 s and $\tau_2$ = 1.41. The accurate estimation of the microscopic rate constants (i.e., the elements of the transition rate matrix) crucially depend on an accurate estimation of the amplitudes of the different exponential terms in the correlation curves. Given the low contrast between the two relaxation times (~30% difference), it is thus likely that the deviations of the inferred rate constants originate from inaccuracies of the estimated amplitudes due to the large overlap of the two exponential components. The fit of the correlation curves and the error estimation by the Markov chain Monte Carlo method are shown in Supplementary Method Figure 7.2.

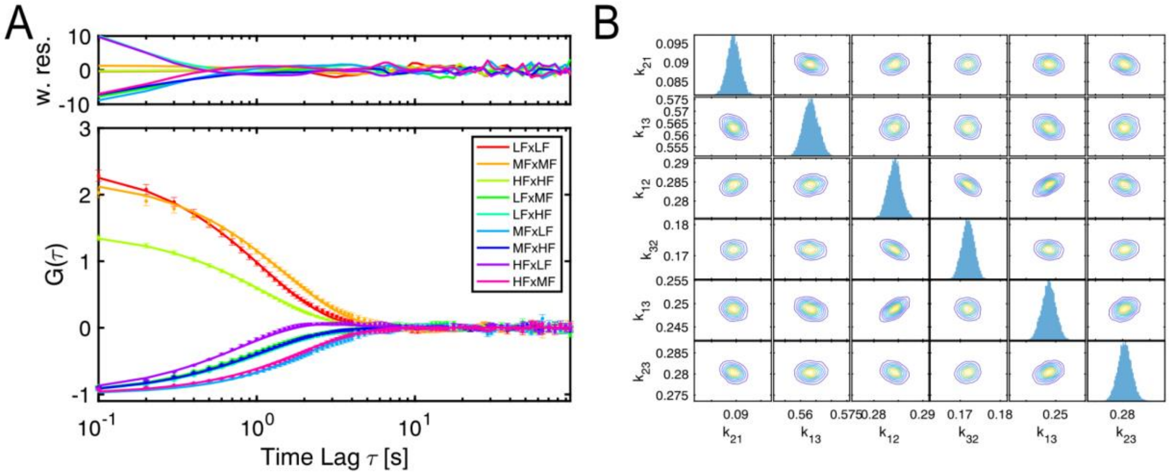


**Supplementary Method Figure 7.2: Correlation analysis of the synthetic dataset of level 2 (three-state system). A)** Filtered-FCS correlation functions (scatter plots) and fits (solid lines). **B)** Posterior distributions of the rate constants of the transition rate matrix. All rates are given in $s^{-1}$.

**Correlation analysis of the degenerate systems (Synthetic Data, Level 3)**

While the FRET efficiency histogram analysis suggested a two-state system for this dataset, we clearly detected multiple relaxation times in the correlation curves that indicated a more complex kinetic network of three or more states. To infer the number of states, we applied the empiric model function with two, three or four relaxation times and compared the BIC values (Supplementary Method Table 7.1 and Supplementary Method Figure 7.3). We found a clear minimum of the BIC for three relaxation times, indicating that the kinetic system involved four states. Based on the existence of two distinguishable FRET states, we speculated that each of these would show a two-fold degeneracy.

While it was not possible to extract the microscopic rate constants reliably, we could compare the relaxation times extracted by the empirical model to the ground truth values. From the filtered-FCS analysis, we estimated relaxation times of $\tau_1$ = 1.3 s (1.1-1.5), $\tau_2$ = 10.8 s (6.6-15.2) and $\tau_3$ = 41 s (30-62) (95% confidence intervals are given in brackets, see Supplementary Method Table 7.1 and Supplementary Method Figure 7.3), which correspond well to the relaxation times determined for the ground truth transition rate matrix of $\tau_1$ = 1.05 s, $\tau_2$ = 8.23 s and $\tau_3$ = 27.00 s. This indicates that correlation analysis can still be used to quantitatively assess the relaxation times corresponding to the transition rate matrix even for complex cases.

**Artifacts in correlation analysis of experimental data**

For experimental data, several artifacts can potentially affect the correlation analysis. Here, we briefly review the most common problems, their effect on the correlation function and how they can be avoided.

*1. Model selection and user bias*

A major advantage of the correlation analysis is that the computation of the correlation function from the signal intensities is free of user bias. However, the quantitative analysis of the obtained correlation functions crucially requires estimating the number of kinetic states (and their FRET efficiencies) to select the correct model function. If the FRET efficiencies of the different kinetic states are sufficiently different, the number of states can be inferred from the framewise FRET efficiency histogram. Importantly, also the number of relaxation times found in the correlation curves (e.g., by use of the empirical model function in conjunction with the BIC) informs on the number of states, where *N* states will result in *N-1* relaxation times. This information was used for the analysis of the synthetic dataset of level 3 with degenerate FRET states. While here the FRET efficiency histogram suggested a two-state system, the correlation curves indicated three relaxation times, in agreement with the four kinetic states of the simulated system. For the filtered-FCS analysis described here, the choice of the number of FRET states is also relevant for the computation of the filtered correlation curves, specifically for the thresholding step for the digitization of the FRET efficiency trajectory. This approach is also inapplicable to degenerate systems containing different kinetic states with identical FRET efficiencies.

*2. Signal spikes due to impurities*

Some of the experimental datasets used in this study showed clear signal spikes both in the donor and acceptor channels that exceed the variation expected for Poissonian noise and do not show the characteristic anti-correlated behavior expected for FRET dynamics (see Fig. 5a,d,g of the main text). Such signal spikes could be of photophysical nature or originate from diffusing impurities (such as unreacted fluorophores) that transiently enter the observation volume. The expected effect is the appearance of an additional fast component in the autocorrelation functions of the donor and acceptor channels for the FRET-FCS analysis. Notably, such signal fluctuations should not affect the cross-correlation function as they should be uncorrelated between the donor and acceptor channels. The effect on the filtered-FCS analysis is more difficult to assess as it depends on whether the step-finding algorithm will erroneously detect the fast changes of the FRET efficiency due to the signal spikes as a transition.

*3. Trace-by-trace variability*

It is generally assumed that all molecules that are considered for the analysis behave identically, however some trace-by-trace variation might occur due to incomplete filtering of dysfunctional molecules, fluorescent impurities, or biologically relevant functional heterogeneity (e.g., originating from a variation of post-translational modifications or allosteric control). Such heterogeneity will skew the average correlation function away from that of the pure species of interest and result in large variations of the shape of the correlation functions. In turn, the experimental uncertainty from the bootstrap procedure will be overestimated, and consequently the reduced chi-squared estimator will be systematically underestimated.

*4. Slow intensity fluctuations*

Slow intensity fluctuations in measurements of immobilized molecules might originate from instabilities of the excitation laser, resulting in slow power fluctuation on the minute to hour timescale, or focal drift due to z-drift of the sample, reducing the detectable signal as the molecule moves outside of the focal plane. As the FRET-FCS analysis is applied directly to the fluorescence intensities, such slow fluctuations will be reflected in the resulting FCS curves. On the other hand, for the filtered-FCS curves the digitized state trajectories are determined based on the FRET efficiency trace which remains unaffected by the signal fluctuations, assuming that the donor and acceptor channels are equally affected and neglecting a change of the noise of the FRET efficiency trace due to the intensity modulation. Slow intensity fluctuation should thus have a minor effect on the filtered-FCS curves.

*5. Photophysical artifacts*

Correlation analysis is sensitive to all effects that modulate the fluorescence intensity. Common unwanted photophysical effects are photoblinking, e.g., due to the population of long-lived dark states such as radical ions[25], and intensity changes due to spectral shifts[26]. Special care should be taken in correlation analysis to avoid that photoblinking of the acceptor fluorophore is interpreted as FRET dynamics, which will feature as a prominent term in the cross-correlation function. Photoblinking of the donor fluorophore is readily detected from a drop of the intensity to the background level. Photoblinking of the acceptor can be identified by the use of alternating laser excitation schemes that include the intermittent probing of the acceptor fluorophore[27,28], allowing to exclude those time intervals where the acceptor was inactive. Generally, the formation of long-lived dark states can be reduced by the application of reducing-oxidizing systems, e.g. by the addition of TROLOX[25].

**Supplementary Method Table 7.1: Analysis results of the correlation analysis.** Rate constants are reported as fitted value ± 95% confidence interval. All rate constants are given in $s^{-1}$ and relaxation times in s. States are ordered from low to high FRET efficiency, i.e. 1=low-FRET, 2=mid-FRET, 3=high-FRET. For asymmetric confidence intervals (dataset 3, empirical three-exp. model), the lower and upper bounds of the 95% confidence intervals are given in brackets.

| **Dataset** | **Method** | $k_{12}$ **[s$^{-1}$]** | $k_{21}$ **[s$^{-1}$]** | $k_{13}$ **[s$^{-1}$]** | $k_{31}$ **[s$^{-1}$]** | $k_{23}$ **[s$^{-1}$]** | $k_{32}$ **[s$^{-1}$]** | $\chi_r^2$ |
|---|---|---|---|---|---|---|---|---|
| **1** | FRET-FCS | 0.142 ± 0.002 | 0.210 ± 0.002 | - | - | - | - | 1.07 |
| | fFCS | 0.142 ± 0.002 | 0.212 ± 0.002 | - | - | - | - | 0.998 |
| **2** | fFCS | 0.091 ± 0.003 | 0.284 ± 0.003 | 0.560 ± 0.005 | 0.249 ± 0.003 | 0.173 ± 0.003 | 0.280 ± 0.003 | 1.83 |
| | | $\tau_1$ **[s]** | $\tau_2$ **[s]** | $\tau_3$ **[s]** | $\tau_4$ **[s]** | | **BIC** | $\chi_r^2$ |
| **3** | empirical. 2 exp | 1.6 ±0.2 | 23.4 ± 1.2 | - | - | - | 71.38 | 0.13 |
| | empirical, 3 exp | 1.3 (1.1-1.5) | 10.8 (6.6-15.2) | 41 (30-62) | - | - | 68.89 | 0.03 |
| | empirical, 4 exp | 1.3 (1.1-1.5). | 8.2 (6.3-16.1) | 11.2 (6.2-16.8) | 40 (30-55) | - | 91.19 | 0.03 |

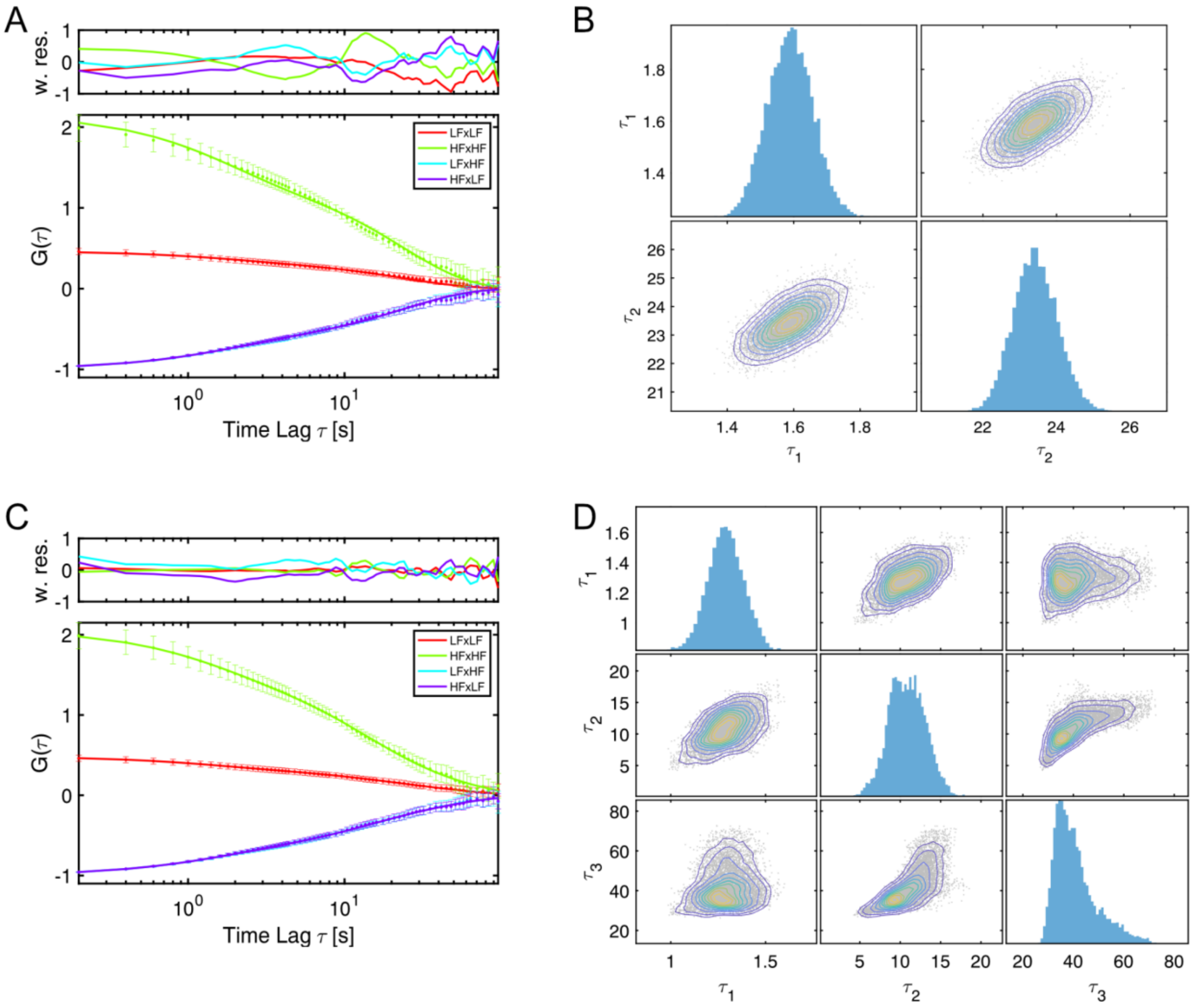


**Supplementary Method Figure 7.3: Empirical analysis of the correlation functions for synthetic dataset of level 3 involving degenerate FRET states.** The data were analyzed using two exponential terms (A-B) or three exponential terms (C-D). **A,C)** Filtered-FCS correlation functions (scatter plots) and fits (solid lines). **B,D)** Posterior distributions of the relaxation times. All times are given in seconds.The $\chi_r^2$ changes from 0.13 to 0.03 and the BIC decreases from 71.38 to 68.89, indicating that the three-exponential model is to be preferred (see Supplementary Method Table 7.1). Inclusion of a fourth component results in an increase of the BIC to 91.19 and shows no further decrease of the $\chi_r^2$ (data not shown).

**Supplementary Method 8: Edge finding (CK)**

Our method is based on the Chung-Kennedy filtering approach previously used in ion channel experiments[29] and subsequently by Haran in single molecule FRET experiments[30]. Applied to a time-series of points representing donor intensity, acceptor intensity or FRET ratio, we associated each data point with a 'forward' window containing a fixed number of later points and a 'backward' window a number of earlier points. The standard deviation (or variance) of the points in such a window will be largest when an edge occurs near the center of the window. By monitoring the standard deviation of the forward and backward windows when scanning through the data time trace, the location of transition edges can be identified by the maxima of these standard deviations (or variance)[31]. In our implementation for the data in this study, we considered only the FRET efficiency time record and constructed only a single window of points. Any window whose standard deviation exceeded a threshold and was also a local maximum was identified to contain a transition edge at its center data point. The window size and critical standard deviation value were determined empirically from the training sets provided with the challenge and verified by inspection by an experienced user. Next, the forward and backward windows were constructed around these potential transition edge locations and the two sample t-test was applied. Transition edges were accepted if the significance level (alpha) of the t-test comparing these data windows examining the FRET states before and after the potential transition edge was above an empirically determined value set by examining the training sets. For the 2 level systems, parameters were: Local window size = 5 data points; threshold for FRET efficiency variance in the window = 0.03; t-test alpha = 0.6. For the 3 level system: Local window size = 5 data points; threshold for FRET efficiency variance in the window = 0.018. t-test alpha = 0.4. Bleaching and blinking events can be removed in a pre-processing step by identifying segments where the sum of donor and acceptor intensities are below a threshold. Removal of blinking and bleaching was only performed on the experimental data but not on the simulated data where the effect was absent. Events immediately before or after removed intervals were not used for kinetic analyses.

Although not required here, it is notable that for more challenging data that Haran has demonstrated increased sensitivity in transition edge detection is possible by exploiting the expected anti-correlation of donor and acceptor intensities associated with transitions of FRET efficiency[30]. This method was implemented by examining donor and acceptor intensities separately, forming local 'forward' and 'backward' windows for each, and identifying maxima in the sum of the donor and acceptor variances in these windows to reveal transition edges.

Once transition edges were identified, FRET states were categorized according to Gaussian fitting of all FRET data points. FRET = 0.55 was the dividing line between states for the 2 state systems, and FRET = 0.35 and FRET= 0.7 were the dividing lines between states for the 3 state systems as determined by locating local minima in the histograms of all FRET values. The dwell time in each state between edges was calculated and histograms of dwell times were assembled for each FRET state. Fitting exponential decay functions to the dwell time histograms yielded estimates of the apparent rates of transitions out of the states. Multiplying apparent rates by the fraction of transitions to a specific state divided by the total number of transitions out of a state ('branching ratio') gives the true transition rate in the 3 state system[32,33].

Additional description of this Chung-Kennedy edge detection method is published elsewhere[31]. The source code is available for download at: https://www.physics.ncsu.edu/weninger/KinSoft.html .

**Supplementary Method 9: Edge finding (k-means)**

The k-means edge finding method is based on an iterative clustering algorithm that assigns data into groups based on the similarity of each point group properties[34–37]. Before performing clustering, the data is preprocessed to remove bleaching and blinking. For this project, the simulated data did not require preprocessing, whereas bleaching and blinking was removed from the experimental data by excluding data points where the sum of donor and acceptor intensities was below a threshold determined by inspection of traces (threshold 27 after 22 point smoothing). The events immediately before or after removed intervals were not used for kinetic analyses.

Our application can apply the k-means edge finding algorithm to time records of donor intensity, acceptor intensity, or FRET efficiency, or a 2-dimensional representation of donor and acceptor intensities. In this project, we only considered the FRET efficiency data. The clustering algorithm groups the FRET efficiency data points without consideration of the time aspect of the record. The number of clusters to be formed is selected based on the apparent number of states in a histogram of all FRET efficiency points for all time traces. There are more formal approaches in k-means theory to determine the number of target clusters[38–40], but those were unnecessary with these data.

The goal is to group the FRET efficiency points into clusters representing the distinct states present in the data. The algorithm proceeds generally by choosing initial values as 'centers' for each group or cluster. All the data points are then assigned to the group which they are closest to by some distance measure without regard to their temporal position in the time trace. The centers of each group are then recalculated by averaging the value of all points in that group. The algorithm then proceeds in an iterative manner whereby all data points are reassigned to the groups minimizing their distance from the new center values. This process of recalculating the center and then reassigning points to the nearest group repeats until points no longer change groups.

In the specific application for the data in this project, we used FRET efficiency values as the input for the k-means algorithm. We determined the number of clusters (or number of 'centers') to use by examining time traces and the FRET efficiency histograms assembled from all the data. To choose initial values for the centers of the clusters, all data point values were ordered by FRET efficiency value and then divided into equally sized groups matching the desired number of clusters. A data point was randomly selected from each cluster to serve as the initial "center" for the cluster. Next, each data point was assigned to the cluster for which the 'distance', which is defined as (data value – cluster 'center' value)$^2$, was minimized. Once all data points were assigned to clusters, the centers for each group were recalculated by averaging the values of all of the data points in the group. Finally, the process was repeated with data points being reassigned based on minimizing the distance to the new centers. This process was repeated until the data points no longer changed clusters. The final value of each "center" was interpreted at the FRET efficiency of a distinct state. All of the data points in a cluster were assigned that FRET state. The data points were then interpreted in the temporal order of the time trace yielding a sequence of FRET state vs. time. Data points where the FRET efficiency state changed were determined as transition edges.

The k-means algorithm assigns points to clusters based on their value without reference to their temporal sequence. For this reason, rare assignments of points to incorrect clusters typically result in time records having transitions to different states that last only one time bin and then return to the previous state. If the data acquisition rate is substantially faster than the rates characterizing the kinetic system, such one time bin events are expected to be rare. For example, if $k_{\text{data acquisition}}$ = 10* $K_{\text{characteristic kinetics}}$ then ~10% of events are expected to be one time bin ; if $k_{\text{data acquisition}}$ = 100*$K_{\text{characteristic kinetics}}$ then ~1% of events are expected to be one time bin . If an erroneous one time bin event breaks up a longer dwell into two shorter dwells, it can have a negative impact on determining the kinetic rates of the system. Therefore, we designed a protocol to remove one time bin events that met certain criteria.

For isolated transitions to other states that last one time bin where the clusters on either side are different from each other, we examined the donor and acceptor values in the clusters to decide how to reassign the one frame. We calculated the squared differences of the donor value on the one time bin event from the averaged donor values of all points assigned to the FRET state of each adjacent cluster and a similar squared differences for the acceptor values:

$$ClosenessToXCluster = \sqrt{(donor\ value - average\ donor\ value\ for\ cluster\ X)^2} + \sqrt{(acceptor\ value - average\ acceptor\ value\ for\ cluster\ X)^2}$$

where X indicates which cluster. The one time bin event was reassigned to the cluster for which the *ClosenessToXCluster* was smallest. If the clusters on either side of a one time bin transition were the same, we used an average of the donor and acceptor values in the points in that cluster for the rest of that particular trace to calculate the closeness to the current cluster *ClosenessToCurrentCluster* and an average of the points in the sections immediately adjacent to the one bin event to calculate closeness to the adjacent cluster *ClosenessToAdjacentCluster* . We then calculated *DifferenceInDistances* = abs(*ClosenessToCurrentCluster -ClosenessToAdjacentCluster )* and the standard deviation of points in the adjacent cluster *stdAdjacentCluster = standard Deviation of all donor (or acceptor) values of adjacent cluster* . If the *ClosenessToAdjacentCluster* < *ClosenessToCurrentCluster* and the *DifferenceInDistances* < *stdAdjacentCluster* for both donor and acceptor signals, then we assigned the point to the state of the adjacent cluster. Otherwise, we left it in the original cluster, maintaining an event lasting only one time bin.

When multiple one time bin events occurred sequentially, we averaged the FRET efficiency in the 3 or more one time bin events and assigned all of them to the same cluster that was closest.

Once all points were finalized into clusters, the dwell time of the behavior in a state was measured as the time between edges and assembled into histograms for all events in a given state (first and last events in every trace as well as events preceding and following blinking were discarded). The histograms were fit with exponential decay functions to determine the lifetimes. For the 3 state system, the true transition rates were determined from the apparent rates by multiplying by the fraction of transitions to a specific state divided by the total number of transitions out of a state (‘branching ratio’)[32,33].

The source code is available for download at: https://www.physics.ncsu.edu/weninger/KinSoft.html .

**Supplementary Method 10: Step finding**

***Step Finding Software***

Analysis with step finding was performed using an in-house implementation of a step identification algorithm written in Python (https://github.com/SMB-Lab/PyStepFinder) that sorts one-dimensional data into piecewise line segments without enforcing a number of states or kinetic network. Sorting is accomplished through iterative forward addition of line segments to a piece-wise linear fit of each trace. At each iteration, starting with one line for the whole trace, all current line segments are tested to find the optimal position at which to split the segments into two segments. The sum of squared residuals ($X^2 = (x - \underline{x})$, with $x$ the value of the test statistic and $\underline{x}$ the mean of the test statistic for the segment) is used as the quality of fit statistic for the fit of the two new segments. This value is compared to the previous segment and the addition of a segment is accepted if the improvement in the quality of fit is significant. This process is summarized visually in (Supplementary Method Figure 10.1). Several options can be specified in performing step finding with this implementation, including choice of the statistic of interest for fitting (mean, rms, variance, slope), a sliding window size for calculating running averages of observed statistics of interest for comparison to the segment means, a minimum length of each line segment (in number of data points), and a threshold for improvement in quality of fit per segment addition as well as a thresholding mode (ratiometric, flat improvement, statistical tests, etc.). For this study, all analysis was performed using the segment means as the statistic of interest with no running average, a minimum segment length of two data points, and the quality of fit threshold set to the variance of each trace. For other test statistics, represents a running average value of the test statistic, in which case the minimum segment length must be at least the size of the averaging window. PyStepFinder is appropriate for other series data as well. We have used a similar tool available in MATLAB previously for analysis of time-series force data from optical tweezer experiments[41]. Currently, PyStepFinder is unable to identify states with degenerate mean signals unless those states are distinguishable by other parameters, such as signal variance. Thus, for this study the step finding algorithm was only used to resolve the non-degenerate FRET efficiencies in degenerate datasets.

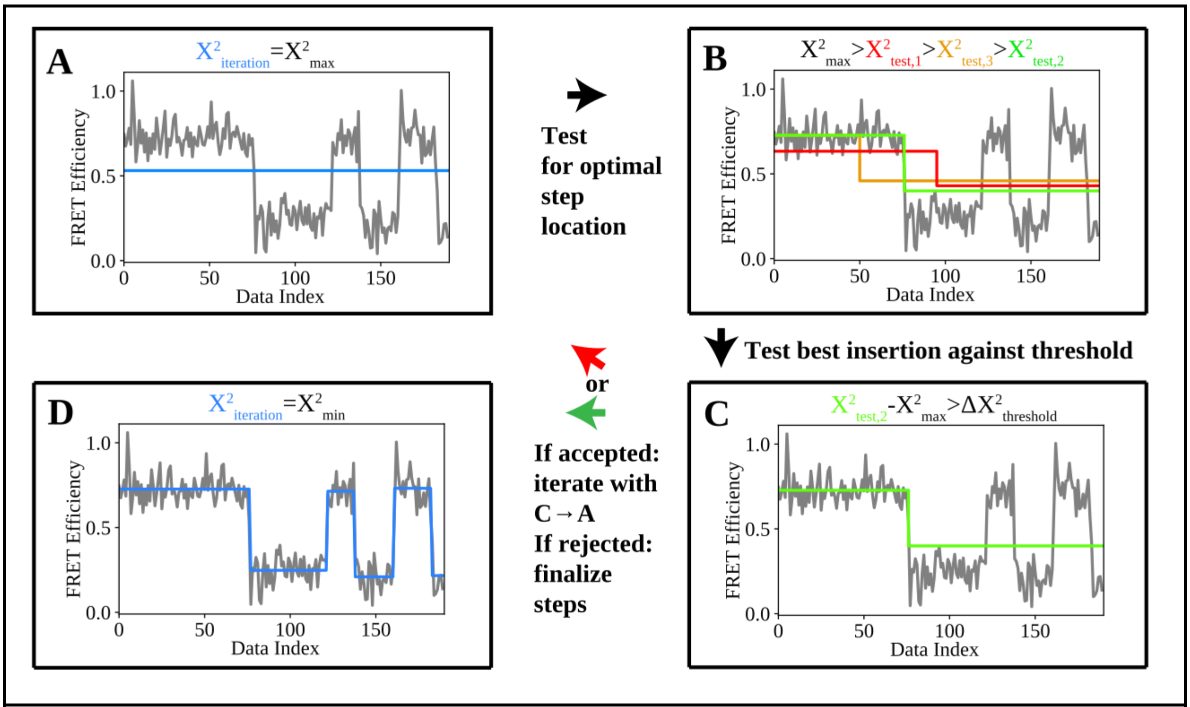


**Supplementary Method Figure 10.1:** Forward addition of steps in PyStepFinder. A) Data is initially fit with a horizontal line at the mean of the data test statistic and initial $X^2$ is calculated. B) Candidate locations are tested for insertion of a step between two segments to replace the original segment. $X^2$ is calculated for each potential replacement and the one with the lowest $X^2$ is taken as the best candidate. C) The change in the $X^2$ calculated for the insertion candidate relative to the original iteration $X^2$ is compared to a threshold value. The criterion for the flat improvement is shown, but other thresholding modes can be defined. If the improvement in $X^2$ satisfies the threshold, iteration continues from A for all existing segments. If the criterion is not met, the segment is accepted as final until no further insertions are accepted for any segment. D) The finalized trace for which insertion of any additional segments would not significantly improve the fit.

***Treatment of Data***

Step finding was performed for each FRET efficiency trace to identify. When FRET efficiency was not provided directly, it was calculated using $Efficiency = I_A/(I_A + I_D)$, where $I_A$ and $I_D$ are the donor and acceptor signal intensities, respectively. Following identification of each transition step in the dataset, each line segment was classified as belonging to one of the non-degenerate FRET states. The number of states was determined by user input following histogramming of FRET efficiency data. The means and widths of these states were determined using Gaussian fits. Line segments then were classified into one of the states based on their mean FRET efficiencies. Dwell times for each line segment were determined by multiplying the number of points in each by the binning resolution of the input dataset. Kinetic rate constants were determined per state from $K_i = 1/\underline{\tau}$, with $\underline{\tau}$ the average dwell time within a state. Rate constants for individual kinetic pathways (i.e., state i to j, $k_{ij}$) were determined by multiplying $K_i$ by the ratio of transitions from state i to state j the total number of transitions from state i, $k_{ij} = K_i \cdot N_{ij}/N_i$. Uncertainty estimates for $k_{ij}$ were determined by calculating the standard error of the mean (SEM) associated with the mean dwell time ($SEM_{\tau,i} = \sigma_{\tau,i}/\sqrt{N_i}$) with $\sigma_{\tau,i}$ the standard deviation of the dwell times. The SEM was then propagated through to each kinetic rate using the standard uncertainty propagation formula, $\Delta k_{ij} = \sqrt{SEM_{\tau,i}^2 \cdot (\partial k_{ij}/\partial \tau_i)^2}$. The first and last line segment for each trace were ignored for this analysis to avoid artificially reduced dwell times associated with each state.

**Supplementary Method 11: STaSI**

Overview of STaSI:

The Step Transition and State Identification (STaSI) method was introduced in 2014 to determine the number of states and step transitions between states for piecewise constant data with a minimum description length (MDL) as the objective function. The step transitions are detected using the Student's t-test and the segments are grouped into states by hierarchical clustering. The optimum number of states is then established using a minimum description length equation that sums the goodness of fit measured using L1 norm and the complexity of the fitting model derived to consider the sparseness of the states and transitions among states. More details on this can be found in the original report by Shuang *et al*. (ref[42]).

Benefits of STaSI include not requiring time-tagged photon counting or photon counting in general. STaSI provides a better resolution to interpret noisy data with fast dynamics to avoid the need for binning. Binning can introduce artifact states in between real states and limits the temporal resolution of single-molecule FRET. STaSI also is objective, requiring no assumptions about the model and no user inputs other than the FRET efficiency trace in determining the number of states and transitions. Finally, the algorithm is written for smFRET data, but could be used for any piecewise constant signal.

Here, we detail how STaSI was applied to the kinSoftChallenge data. While STaSI was developed only for state and step transition identification, we analyze the kinetics of the transitions identified by STaSI with a simple exponential fitting method to maintain its ease-of-use. Overall, STaSI is a user-friendly, objective method developed to analyze a variety of piecewise constant data conditions. STaSI will require future development to handle data with degenerate kinetics and other experimental complexities where quantifying kinetics requires more extensive user-input beyond the scope of the original STaSI report.

STaSI Methods:

The STaSI GUI (Graphical User Interface) determined the state assignments and transitions for all traces in each dataset level (https://github.com/LandesLab/STaSI). The provided donor intensity ($I_D$) and acceptor intensity ($I_A$) values were converted to a FRET efficiency ($E$) using:

$$E = I_A/(I_D+I_A) \tag{11.1}$$

in a 1D vector format compatible with STaSI. The frame rate was noted separately. The STaSI GUI was executed on the FRET E. All traces were analyzed together, as the STaSI GUI is able to concatenate the vectors into one dataset. STaSI outputted the calculated grouped states and step transitions for 1-30 number of states and the minimum description length (MDL) based on the complexity of the model and goodness of fit was calculated for each number of states. The grouped states and step transitions for the global minimum MDL was saved as a FRET E vector as the output of STaSI. Non-physical states identified with FRET E > 1 or < 0 were not considered in the final number of states or any of the kinetic analysis. The FRET E was the assigned state levels of the STaSI output. The sigma FRET/σ(FRET E) was calculated from the standard deviation of the raw FRET E data assigned to each state level in STaSI.

The total duration of inference for the STaSI analysis was recorded using the tic and toc functions in MATLAB R2018a executed after the user input. All the computations were done using a Dell desktop computer with an Intel(R) Core(TM) i7-8700 CPU @ 3.20 GHz processor, 16.0 GB RAM and 64-bit Windows 10 Enterprise (2018) operating system.

The kinetics were calculated by an exponential fit of the cumulative distribution of dwell times spent in each state. The original STaSI report only indicates FRET state levels and the step transitions between levels. Thus a separate MATLAB script for kinetic analysis was written similar to previous work in literature by Landes *et al*. (ref[43]) and Benitez *et al*. (refs[44,45]). The STaSI output vector was converted to dwell times in each state using the state level, indices of the transitions, and the frame rate. Any false transitions caused by concatenating the individual traces together in the STaSI analysis were removed (*i.e.* connecting end of trace i with the beginning of trace i+1). The cumulative probability distributions of dwell times for each state were fit to a simple exponential decay model,

$$P(t > \tau) = \sum_{i=1}^{n} A_i \, e^{-k_i \tau} \qquad (11.2)$$

where $P(t>\tau)$ is the cumulative distribution of observing a dwell time of $\geq\tau$ for a given time $t$, $A$ is the amplitude, $k$ is the rate, and $n$ being the number of components[46]. MATLAB's built in 'fit' function using the Trust-Region-Reflective Least Squares was used with the inverse of the mean value of dwell time as an initial guess for $k$ and 1 for $A$.

For data that resulted in more complexity than simple two-state transitions (Figures 3-5 of the main text), we further analyzed the resulting values from Equation 11.2 by[44,47]:

$$k_{ab} = A_{ab} / \left[\sum_{i=1}^{n} \left(\frac{A_{ab}}{k_{ab}}\right)\right]. \qquad (11.3)$$

Here, $a$ is the starting state and $b$ is the ending state for the kinetic rate transition of interest and $n$ represents the total number of states. $A_{ab}$ can either be extracted from the fit from non-normalized cumulative distributions, or, as we used here, the total number of transitions observed from state $a$ to state $b$. For example, Equation 11.3 states that the rate of state 1 to state 2 transition ($a$=1 and $b$=2) is calculated as the ratio between the number of transitions from state 1 to state 2 divided by the total dwell time in state 1 spent before transitioning any other state (2, 3, … n). This method is used because, $k_{ab} \neq 1/\tau_{ab}$ for non unimolecular reactions[47].

The reported kinetic rate model values were the resulting $k$ from Eqs. 11.2 and 11.3. The uncertainties of rate models were interpreted as the 95% confidence of the $k$ fit results using MATLAB direct syntax, *confint*. For three-state and higher systems, the appropriate error propagation was also carried out to calculate the 95% confidence intervals.

**Supplementary Method 12 & 13: MASH-FRET**

#### a. Determination of the FRET state configuration

The procedure used in the main article to determine the FRET state configuration with the MASH-FRET (bootstrap) and (prob.) methods is based on the conclusions of a preliminary comparative study of several algorithms[48].

The provided acceptor $I_A(t)$ and donor $I_D(t)$ intensity signals were not further processed as they were provided free of background and dye photobleaching. Additionally, the necessary data to correct the differences between donor and acceptor quantum yields, *i.e.,* the control acceptor signal upon acceptor direct excitation, was not part of the provided data sets. FRET-time traces $FRET(t)$ were therefore directly calculated according to:

$$FRET(t) = \frac{I_A(t)}{I_A(t)+I_D(t)}. \quad (12.1)$$

Aberrant FRET values below -0.2 and above 1.2 FRET units were ignored in the following analysis.

Individual FRET-time traces were discretized into FRET state sequences using the algorithm STaSI[42]. Because this algorithm does not make any assumptions about the kinetics of state transitions, and thus prevents the detection of false transitions towards noise-induced artefactual states, it has proven to be the most suitable to identify the genuine FRET states[48]. The maximum number of states to be found in each FRET trajectory was arbitrarily set to a large number, *e. g.* 10.

To group the FRET states of all sequences into one global state configuration, we chose to sort them drawbacks of a one-dimensional distribution, *i.e.*, the merging of state populations having similar FRET values, by splitting the population along an additional axis: the FRET state forwarding the transition in the trajectory. After smoothing with a Gaussian filter, the TDP was modelled with a mixture of isotropic 2D-Gaussians, which centers were locked on a $V$-by-$V$ grid, with $V$ the number of global FRET states. In addition, Gaussian clusters on the TDP diagonal were used to group, and then exclude from the analysis, the artefactual and noise-induced low-amplitude state transitions. To determine the most sufficient model size $V_{opt}$, Gaussian matrices with increasing dimension, *i.e.* $V$ =2 to 10, were fitted to the TDP using an expectation-maximization (EM) approach, and the model rendering the lowest Bayesian information criterion (BIC), calculated as

$$BIC(V) = (2V^2 + V - 1)log(M) - 2log[l(V)], \quad (12.2)$$

where $M$ is the total number of states in the trajectories and $\ell$ the likelihood of the model, was selected.

With the number of observable FRET states at hand, FRET-time traces were re-discretized into more accurate state sequences with the Bayesian-based algorithm vbFRET. Indeed, although a model-free algorithm provides a better global view of the state configuration, it is not suitable for detecting short-lived states. However, vbFRET is designed for Gaussian-distributed trajectory noise and fails to properly characterize *e. g.* low-photon-count trajectories that generate Poisson noise. In such cases, the sequences generated by STaSI were used. The vbFRET algorithm was constrained to found $V_{opt}$ states at maximum and its well-known propensity to detect artefactual blur states[49] was post-corrected by ignoring all one-data-point states found in trajectories.

To accurately determine the global FRET states, a mixture of multivariate Gaussians, which centers were locked on a $V_{opt}$-by-$V_{opt}$ grid, was fitted to the new TDP using the EM approach mentioned above. Global FRET values were derived from the Gaussian means and the associated errors, $\delta_{FRET}$, from the average Gaussian standard deviations in the x-direction $\sigma_x$, such as:

$$\delta_{FRET,v} = \frac{1}{V-1}\sum \sigma_{x,vi}. \tag{12.3}$$

#### b. Estimation of the transition rate constants

Transition rate constants were determined in two different ways: the *bootstrap* and the *probabilistic* (*prob.*) approaches. The *bootstrap* approach is only suitable for non-degenerate state systems, *i.e.*, for states with distinct FRET values, whereas the *prob.* approach suits all types of systems.

##### The "bootstrap" approach

Dwell times $\Delta t$ associated to each global FRET state were collected from accurate FRET state sequences and normalized cumulative distributions $F$ were built. The complementary distribution was subsequently fit with a single exponential function such as

$$1 - F(\Delta t_v) \sim exp\left(\frac{-\Delta t_v}{\tau_v}\right), \tag{12.4}$$

where $\tau_v$ is the state lifetime. As time-binned data suffer from the absence of very short dwell times, the normalized complementary cumulative histogram $1 - F(\Delta t_v)$ of dwell times $\Delta t_v$ is used instead of raw counts. This minimizes the impact of the first histogram bins while preserving the overall shape.

Transition rate constants $k$ were derived from the state lifetimes and the numbers of transitions $w$ using the relation

$$k_{v,v\prime} = \frac{w_{v,v\prime}}{\tau_v \sum w_{v,k}}. \tag{12.5}$$

Please note that for a two-state system, the transition rates are the direct inverse of the lifetimes, *i.e.*, $k'_{v,v\prime} = \frac{1}{\tau_v}$.

The outcome of such analysis are single estimates of the rate constants. To estimate the error $\delta_k$ on rate constants $k$, the variability of state lifetimes across the trajectory sample is evaluated using the bootstrap-based analysis called BOBA-FRET[50]. BOBA-FRET infers the bootstrap means and bootstrap standard deviations of all fitting parameters for the given sample, including $\tau$. The variability can then be propagated to $k$ such as:

$$\delta_{k,v,v'} = \frac{\sigma_{\tau,v}}{\bar{\tau}_v}\bar{k}_{v,v'} \tag{12.6}$$

where $\bar{\tau}$ and $\sigma_\tau$ are respectively the bootstrap mean and standard deviation of parameter $\tau$ and $\bar{k}$ is the rate constant derived from $\bar{\tau}$ using Eq. 12.5. Intervals with 95% confidence were given as $\bar{k} \pm 2\delta_k$.

**The "probabilistic" approach**

The presence of degenerate states usually breaks the single exponential shape of the dwell time distribution, resulting in sums and convolutions of multiple distributions. The *probabilistic* approach first solves the state degeneracies, *i.e.* the numbers of degenerate states hidden behind the same FRET values, from the shapes of ensemble dwell time distributions, and second, optimizes the transition probability matrix for the set of FRET state sequences.

Phase-type distributions (PH) are used, *e. g.* in queuing and insurance risk theory, to estimate the time, $t_{abs}$, a Markov jump process takes to reach an absorbing state, depending on the number of phases $D$ it can go through. Such a jump process involving 3 phases is illustrated below (Supplementary Method Figure 13.1)

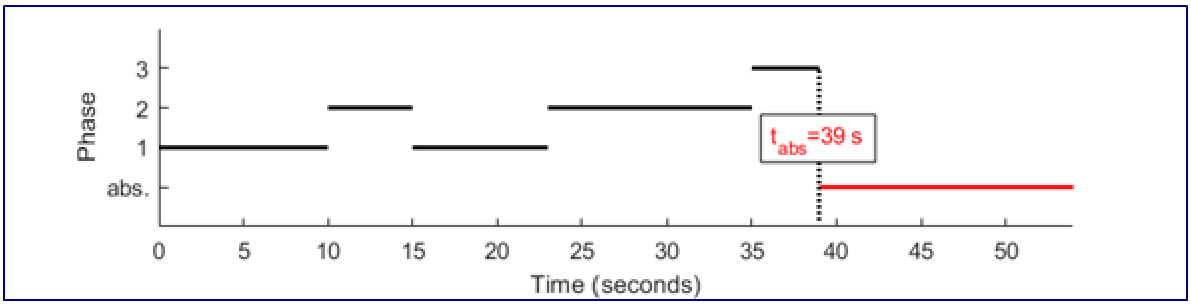


**Supplementary Method Figure 13.1.** Illustration of a Markovian jump process. going through a number of phases $D = 1\ldots 3$, i.e. the number of degenerated states, before reaching the absorbing state, thus, the observable state transition, i.e. between two observed FRET states.

In comparison to our problem, the phases labeled 1 to $D$ are the degenerate states behind a same FRET value, the Markov jump process characterizes the transitions between these degenerate states, the absorbing state is any state having a different FRET value, and the absorbing times $t_{abs}$ are the dwell times $\Delta t$ measured in the state sequences. Therefore, PH distributions make perfect candidates to model the dwell time histograms compiled for a degenerate state system. As the data provided for analysis were time-binned trajectories, discrete PH distributions (DPH) were used instead. The DPH probability density function $f$ depends on transition probabilities between degenerate states and to the absorbing state (state 0), $p$, as well as on starting probabilities, $\pi$. All in all, it is expressed as:

$$f(\Delta t_v) = (\pi_1, \pi_2, \dots, \pi_D) \times \begin{pmatrix} p_{1,1} & p_{1,2} & \cdots & p_{1,D} \\ p_{2,1} & p_{2,2} & \cdots & p_{2,D} \\ \vdots & \vdots & \ddots & \vdots \\ p_{D,1} & p_{D,2} & \cdots & p_{D,D} \end{pmatrix}^{\Delta t_v - 1} \times \begin{pmatrix} p_{1,0} \\ p_{2,0} \\ \vdots \\ p_{D,0} \end{pmatrix} = \pi T^{\Delta t_v - 1} \mu \quad (13.1)$$

Where $\boldsymbol{\pi}$ is called the initial distribution of phases, $\boldsymbol{T}$ the sub-intensity matrix and $\boldsymbol{\mu}$ the exit rate vector.

After re-binning the dwell times using a bin size 10-time larger than the resolution time in order to minimize the impact of the lack of very short dwell times in time-binned data while preserving the overall shape, dwell time histograms were modelled with a DPH involving $D$ degenerate states. To determine the most sufficient model size $D_{opt}$ for each histogram, DPHs with increasing dimensions, *i.e.* $D$ =1 to 4, were fitted using an EM approach described previously[51] and the combined model rendering the lowest BIC was selected. In our particular case, the BIC of the combined model was calculated as the sum of the BIC values obtained for individual dwell time histograms, such as:

$$BIC = \sum B\,IC(D_v) = \sum np(D_v) \times log(M_v) - 2 \sum l(D_v), \quad (13.2)$$

where $M_v$ is the number of dwell times in the histogram, $\ell$ the likelihood of the model, and where the number of free parameters $np$ is calculated as:

$$np(D) = D^2 - 1 \quad (13.3)$$

With the final model size at hand, we determined the corresponding transition rate constants by applying the Baum-Welch[52] algorithm to state trajectories, i.e., to noiseless trajectories, in which the state assignment is inflexible. Therefore, the algorithm only optimizes the transition probability matrix by iterating expectation and maximization of state probabilities at each time bin of each state trajectory. It eventually converges to a maximum likelihood estimator of transition probabilities that are then converted into rate constants, using the relation

$$k_{j,j'} = \frac{p_{j,j'}}{t_{exp}} \quad (13.4)$$

where $k_{j,j'}$ is the rate constant that governs transitions from state $j$ to state $j'$ (in seconds$^{-1}$) and $t_{exp}$ is the bin time in trajectories (in seconds).

The negative and positive errors $\delta_k^-$ and $\delta_k^+$ on rate coefficients were estimated via a 95% confidence likelihood ratio test described elsewhere[53], giving an estimated range delimited by the lower bound k-$\delta_k^-$ and the upper bound $k$+$\delta_k^+$.

To ensure the validity of the inferred model, a set of synthetic state trajectories is produced using the kinetic model parameters and the experimental mensuration (sample size, trajectory length), which is then compared to the experimental data set[47]. Special attention is given to the shape of each dwell time histogram, the populations of observed states and the number of transitions between observed states.

#### In practice

To run MASH-FRET, you will need a computer equipped with MATLAB and the following toolboxes:

- Symbolic Math Toolbox
- Image Processing Toolbox
- Statistics and Machine Learning Toolbox
- Curve Fitting Toolbox

MASH-FRET was mainly developed on MATLAB2016a (Windows 8.1) but has recently passed to MATLAB2020b. Therefore, we can guarantee proper functioning only for MATLAB2020b on Windows 8.1. Computation times were measured on a computer equipped with an Intel Core i7-3632QM CPU (2.2GHz) and 8GB of RAM.

The steps to reproduce the results obtained in the main article are the following[54]:

1. Install and start MASH-FRET v.1.3.2 as described in the online documentation (https://rna-fretools.github.io/MASH-FRET/Getting_started.html)
2. Go to MASH-FRET's menu *Routines* > *Standard analysis* > *All steps* and select the set of files to analyze
3. A first message box pops up: enter the number of FRET states if known or press "No" otherwise
4. A second message box pops up: choose the proper noise distribution according to your data set
5. Once the analysis routine is completed, you can find the analysis summary in file *[data file name]_results_[J]states.txt* at the same location as your data files.

#### Notes

MASH-FRET delivers transition rates restricted to 2-state systems up to version v.1.2.1 and below. This has been corrected. The development of MASH-FRET (prob.) has been initiated by the kinsoftchallenge to solve degenerate FRET-state systems, i.e., FRET states comprising kinetic heterogeneity. Therefore, the analyses of round 1 and 2 have been repeated with the new software version of MASH-FRET (prob.) and are labelled as "post-ground truth submission" where necessary. Further, the determination of the number of observable FRET states in MASH-FRET was modified meanwhile the submission process, which led to discrepancies between the bootstrap and the prob. method (compare Fig. 5 of the main text). The most recent version v.1.3.2 of the software yields two observable FRET states for all three data sets for both methods and as presented for MASH-FRET (prob.) in Fig. 5b,e and h of the main article.

**Supplementary Method 14: postFRET**

The concept of the postFRET analysis[47] is to fit the experimental single-molecule FRET (smFRET) data first then simulate similar data for comparison. Source codes (MATLAB) are available at https://github.com/nkchenjx/postFRET. A simple thresholding method is used, i.e. set a threshold (e.g. the FRET value in the middle of two states) to distinguish the two states. This kind of analyzed results contains two major errors: (1) state miss-assignment due to the noise, (2) state miss-assignment due to camera blurring. After assigning states with the threshold, >hundreds of virtual data are simulated in the hope that one can find one or more trajectories that look just like the experimental data using the same analysis method, e.g. the thresholding method. Because we know the ground truth of the simulated data, we assume that the hidden truth of the real experimental data is the same as the simulated data that looked the same (minimizing L1-norm, the absolute values of the percentage errors, as the judging standard in Ver 1.0 and 2.0). L1-norm is used instead of L2-norm (such as the least square root method). The former works better in many simulated conditions in postFRET.

The guessing algorithm of the simulated data used in the codes is a semi-exhaustive searching algorithm called JCFit (available on GitHub, https://github.com/nkchenjx/JCFit), a fitting algorithm that searches a parameter in an equation (model) within a defined boundary. The searching spacing is exponentially distributed away from the initial guess to the boundary. E.g. -10 to 10 are the boundaries and 1.0 is the initial guess, and 0.1 is the searching accuracy and ln(2) is the exponential factor, then the searching space is [1.0, 1.1, 1.3, 1.7, 2.5, 4.1, 7.3, 10] going up, and [1.0, 0.9, 0.7, 0.3, -0.5, -2.1, -5.3, -10] going down. The boundaries in version 2.0 are set mobile among searching iterations.

The MATLAB codes are divided into a few steps with the file name Sx_xxxx.m, where Sx represents the step order. Examples are given for the training data level 1.

**Step 1.** Load data. Load all trajectories into one single matrix and mark the end of each trajectory in a separate vector. The photobleaching information is analyzed and used later in the postFRET analysis.

**Step 2.** Load key. If the key of the rate constants is known (as in the training data), load the key (type in manually). This step is not needed for real data. Thus, if the key is not known, give a random guess based on the number of states observed.

For the level 1 training data, the key is given: **[0, 0.666; 1, 0].** Note the diagonal in postFRET is always 0. The direction is column to row, 90 degree rotation of the kinSoftChallenge format.

**Step 3.** Find the noise. Normalize the total counts of the acceptor and donor channel for each trace (**Supplementary Method Figure 14.1**). Then analyze the noise model in the sum of the two channels using the standard deviation of the normalized total signal. Then calculate the average noise of each channel. The latter is used to simulate the trajectory later. The noise model is pretended to be unknown and a Gaussian model is identified for the training data.

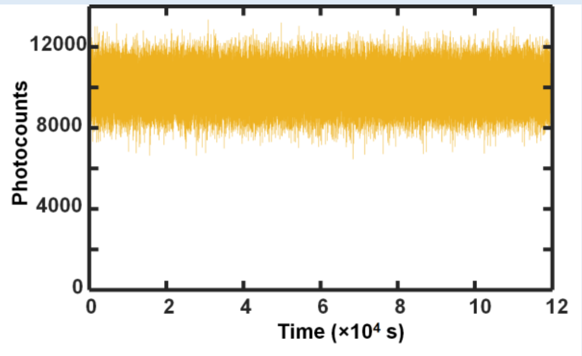


**Supplementary Method Figure 14.1.** The total photocounts of the 100 traces. This trajectory is used to analyze the noise level of the signal.

**Step 4.** Manually determine the number of states (**Supplementary Method Figure 14.2**), the state values and analyze the data using the simple thresholding method (**Supplementary Method Figure 14.3**). The thresholds are set in the middle of two adjacent states.

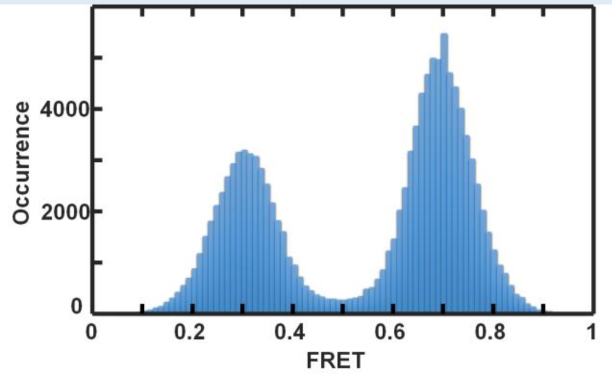


**Supplementary Method Figure 14.2.** Two states are identified with FRET values of 0.3 and 0.7. The uncertainty is the sigma of the two Gaussian peaks.

The thresholding analysis gives the transition rates: **[0,0.63; 0.92, 0]** (**Supplementary Method Figure 14.3**). The detailed procedure has been described in the cited paper and its supporting information. This value is only slightly biased to the truth by the noise because the signal-to-noise level is high in this set of data. It will be more biased with a higher noise level. Briefly[47]

$$k_{if} = \frac{N_{if}}{\sum_{f=1}^{N} t_{if}} = \frac{N_{if}}{t_{if}}$$

Where $k_{if}$ is the rate constant from state $i$ to state $f$, $N_{if}$ is the fitted total number of transitions from state $i$ to state $f$, $t_{if}$ is the sum of the dwell times of state $i$ to state $f$, and $t_i$ is the total dwell time in state $i$. If the dwell time is only one pixel, it is merged to the previous state as noise. The first and the last transition of a trace is also removed from the counting to avoid the edge effect.

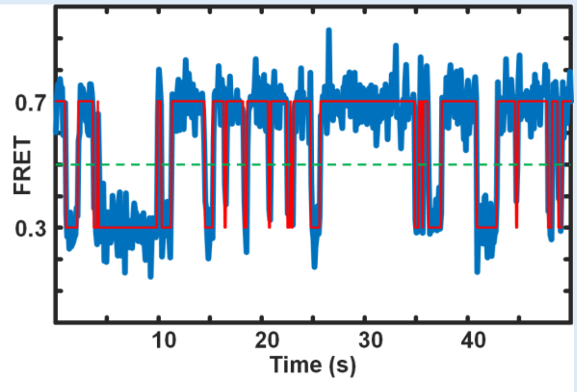


**Supplementary Method Figure 14.3.** Thresholding analysis of the FRET trajectory (showing the first 50 s data). The dashed line indicates the threshold.

The total computational time of all the above steps is negligible on a regular desktop computer for this set of data (less than 1 minute). The thresholding state identification takes 1.3 s, which is linearly proportional to the length of the data.

**Step 5.** postFRET analysis by simulating data with the same photoblinking value and the same noise to find similar trajectories to the raw data. The rate constant key is ignored here so one can just compare the rate constants between the raw data and the simulated data. The goal is to minimize the difference by searching the "real rates" of the simulations. The scoring equation is:

$$WL = \sum_{i} \left| \frac{R_{E,i} - R_{S,i}}{R_{E,i}} \right|$$

where $R_E$ is the analyzed rate of the experimental data, and $R_S$ is the rate of the simulated data, $i$ is the $i^{th}$ non-zero rate in the rate matrix.

The searching space (defined by boundaries) is from ½ to 2 times the initial guess that has been exponentially distributed from the guess value to the boundaries. Thus, the boundary is changing when the initial guess changes from iteration to iteration.

For the level 1 data, the algorithm finds a simulated trajectory very similar to the raw data in the first iteration and this is repeated a number of iterations showing the variation of the scores (**Supplementary Method Figure 14.4**). The best score is approaching the theoretical best 0% during some iterations. However, the value varied in a small region from 3% to 0%. Because no consistent score decay is observed, all values of all iterations are kept for error analysis.

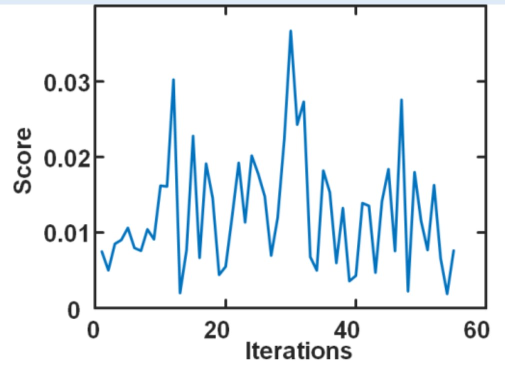


**Supplementary Method Figure 14.4.** The score of the postFRET over the iterations. The computation time of each iteration is ~100 s (single CPU) on a regular desktop with a 3.4 GHz Intel i7 CPU. Parallel computing of multiple CPU and GPU has not been activated.

The mean value and standard deviation of the rate constants of the 55 iterations are **[0, 0.66±0.02; 0.95±0.02, 0]**. Comparing to the key **[0, 0.666; 1, 0]**, this is a better value than the results obtained from the simple thresholding method **[0, 0.63; 0.92, 0]**. An example trajectory of the 55 guesses is shown in **Supplementary Method Figure 14.5**. One can see that the simulated data is different from the raw data but carries similar state and kinetic information.

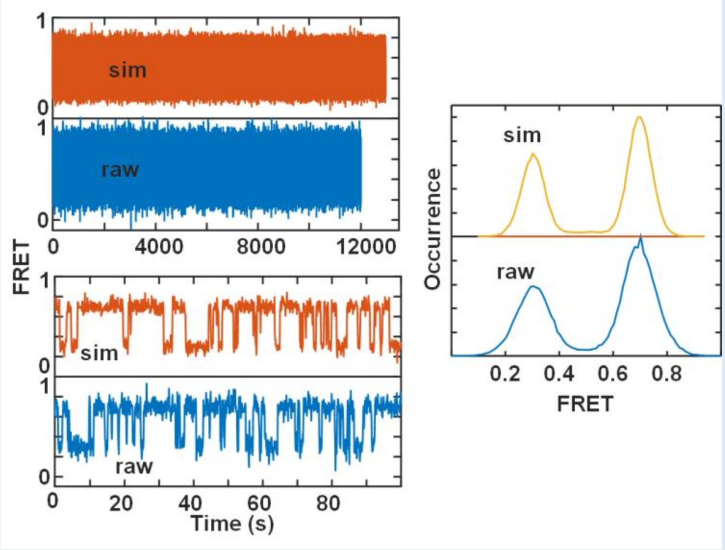


**Supplementary Method Figure 14.5.** An example simulated trajectory comparing to the raw data. The bleaching time is stochastic. The simulated data in this example is longer than the real data. Zoom in on the trajectory of 100 traces showing a similar pattern between the simulated (sim) data and the raw data (left). The distribution of the two states also shows a similar pattern (right). The raw data is wider in the distribution than the simulated data indicates that the noise level is slightly lower-estimated in the simulated data. No attempt is tested to increase the noise level of the simulation to match the raw data.

The error bars associated with the mean rate constants of these iterations are obtained from the standard deviation of the rates among iterations. Two times the error bars represent 95% confidence of the boundaries. The positive (upper) and the negative (lower) standard deviation are slightly different in this set of data which is also reported. Because the noise is relatively symmetric in the data, the difference is small.

**Running time.** The total computational time of steps 1 to 4 is negligible on a regular desktop computer (CPU Intel i7 3.4 GHz) for this set of data (less than 1 minute). The thresholding state identification takes 1.3 s, which is linearly proportional to the length of the data. The running time of step 5 is ~100 s each iteration for the training data, which is linearly proportional to the length of the raw data, and linearly proportional to $n(n-1)$, where $n$ is the number of states since the codes search each transition in an iteration. Parallel and GPU computation can significantly reduce the simulation time for postFRET.

The postFRET code works for two-state and multi-states smFRET analysis but is not coded to detect degenerated states. The code only analyzes FRET values and does not analyze the donor and acceptor channel separately. To make that change, the concept should still work but a significant amount of modification is required.

The postFRET code is expected to be more competitive in analyzing slightly noisier data and data with very fast transition rate constants approaching the time resolution of data collection. That kind of data has significant amounts of camera blurring events. However, it cannot analyze too noisy data when state mis-assignment becomes too large for the thresholding method. For those data, binning must be applied to increase the signal-to-noise ratio or other state-identification methods are needed to replace the thresholding method.

### 4 Supplementary Tables

We provide here all inferred values concerning the data discussed in Figs. 4 and 5 of the main text. This is an excerpt of the complete inferred results found in the Supplementary Datafiles (excel sheets). The inferred FRET efficiency levels and kinetic models are specified using the following nomenclature:

***Nomenclature***

| | | | |
|---|---|---|---|
| FRET E | $E_1$ | $E_2$ | ... |
| kinetic model | 0 | $k_{12}$ | ... |
| | $k_{21}$ | 0 | ... |
| | ... | ... | ... |

**Supplementary Tables 1**

The rate constants of the GT and the inferred models shown in Fig. 4 of the main text. The rate constants are specified in $s^{-1}$. The full submission, including standard deviations of the FRET efficiencies and uncertainties of the rate constants, can be found in the Supplementary Datafiles. Please note: the order of states in the Supplementary Datafiles corresponds to the submission of the participants and may thus differ from the order given here.

***0) Ground truth***

| | | | | |
|---|---|---|---|---|
| FRET E | 0.18 | 0.18 | 0.73 | 0.73 |
| kinetic model | 0 | 0.053 | 0 | 0.018 |
| | 0.080 | 0 | 0.250 | 0 |
| | 0 | 0.680 | 0 | 0 |
| | 0.032 | 0 | 0 | 0 |

***1) Pomegranate***

| | | |
|---|---|---|
| FRET E | 0.181 | 0.714 |
| kinetic model | n.a. | n.a. |
| | n.a. | n.a. |

***2) Tracy (HMM)***

| | | | | |
|---|---|---|---|---|
| FRET E | 0.18 | 0.18 | 0.73 | 0.73 |
| kinetic model | 0 | 0 | 0.0937 | 0.0291 |
| | 0 | 0 | 0 | 0.8846 |
| | 0.953 | 0 | 0 | 0 |
| | 0.0971 | 0.0484 | 0 | 0 |

***3) FRETboard***

| | | | |
|---|---|---|---|
| FRET E | 0.196 | 0.658 | 0.752 |
| kinetic model | 0 | 0.125 | 0.003 |
| | 0.513 | 0 | 0 |
| | 0.027 | 0 | 0 |

##### *4) Hidden-Markury*

| | | | | |
|---|---|---|---|---|
| FRET E | 0.186 | 0.186 | 0.725 | 0.725 |
| kinetic model | 0 | 0.039 | 0 | 0.017 |
| | 0.047 | 0 | 0.246 | 0 |
| | 0.045 | 0.569 | 0 | 0.003 |
| | 0.037 | 0 | 0 | 0 |

##### *5) SMACKS(SS)*

| | | | | |
|---|---|---|---|---|
| FRET E | 0.19 | 0.19 | 0.72542 | 0.72542 |
| kinetic model | 0 | 0.04362695 | 0 | 0.01930565 |
| | 0.063392 | 0 | 0.226308 | 0 |
| | 0 | 0.601065 | 0 | 0 |
| | 0.0351999 | 0 | 0 | 0 |

##### *6) SMACKS*

| | | | | |
|---|---|---|---|---|
| FRET E | 0.19 | 0.19 | 0.71 | 0.71 |
| kinetic model | 0 | 0.0428 | 0.0001 | 0.0195 |
| | 0.0584 | 0 | 0.2254 | 0 |
| | 0.0055 | 0.5985 | 0 | 0.0034 |
| | 0.0359 | 0 | 0 | 0 |

##### *7) Correlation*

| | | | | |
|---|---|---|---|---|
| FRET E | 0.18 | 0.18 | 0.73 | 0.73 |
| kinetic model | n.a. | n.a. | n.a. | n.a. |
| | n.a. | n.a. | n.a. | n.a. |
| | n.a. | n.a. | n.a. | n.a. |
| | n.a. | n.a. | n.a. | n.a. |

##### *8) Edge finding (CK)*

n.a.

##### *9) Edge finding (k-means)*

n.a.

##### *10) Step finding*

| | | |
|---|---|---|
| FRET E | 0.185 | 0.726 |
| kinetic model | 0 | 0.19 |
| | 0.327 | 0 |

##### *11) STaSI*

| | | |
|---|---|---|
| FRET E | 0.2 | 0.7 |
| kinetic model | n.a. | n.a. |
| | n.a. | n.a. |

***12) MASH-FRET (bootstrap)***

| | | | | |
|---|---|---|---|---|
| FRET E | 0.186 | 0.186 | 0.726 | 0.726 |
| kinetic model | n.a. | n.a. | n.a. | n.a. |
| | n.a. | n.a. | n.a. | n.a. |
| | n.a. | n.a. | n.a. | n.a. |
| | n.a. | n.a. | n.a. | n.a. |

***13) MASH-FRET (probabilistic)***

| | | | | |
|---|---|---|---|---|
| FRET E | 0.181 | 0.181 | 0.708 | 0.708 |
| kinetic model | 0 | 0.045 | 0 | 0.024 |
| | 0.050 | 0 | 0.233 | 0 |
| | 0.033 | 0.569 | 0 | 0 |
| | 0.043 | 0 | 0 | 0 |

***14) postFRET***

| | | |
|---|---|---|
| FRET E | 0.19 | 0.73 |
| kinetic model | 0 | 0.1 |
| | 0.139 | 0 |

**Supplementary Tables 2**

Kinetic models for the data shown in Fig. 5a-c of the main text, inferred by the participating groups with the specified tools. Units of the rate constants are in $s^{-1}$. The full submission, including standard deviations of the FRET efficiencies and uncertainties of the rate constants, can be found in the Supplementary Datafiles.

***1) Pomegranate***

| | | | | |
|---|---|---|---|---|
| FRET E | 0.205 | 0.489 | 0.719 | 0.927 |
| kinetic model | 0 | 0.5428 | 0.7998 | 0.6016 |
| | 2.6295 | 0 | 2.8355 | 3.7314 |
| | 1.2125 | 0.9753 | 0 | 1.2084 |
| | 0.9096 | 0.8069 | 0.7324 | 0 |

***2) Tracy (HMM)***

| | | | |
|---|---|---|---|
| FRET E | 0.23 | 0.76 | 0.9 |
| kinetic model | 0 | 0 | 0.03 |
| | 0.9 | 0 | 0 |
| | 0 | 0.029 | 0 |

***3) FRETboard***

| | | | | |
|---|---|---|---|---|
| FRET E | 0.229 | 0.385 | 0.648 | 0.842 |
| kinetic model | 0 | 0.1341376 | 0.1197335 | 0.1764494 |
| | 0.5357143 | 0 | 0.1897321 | 0.1636905 |
| | 0.2962113 | 0.0849598 | 0 | 0.2870264 |
| | 0.1867587 | 0.0371747 | 0.1079837 | 0 |

***4) Hidden-Markury***

| | | |
|---|---|---|
| FRET E | 0.222 | 0.802 |
| kinetic model | 0 | 0.383 |
| | 0.36 | 0 |

***5) SMACKS(SS)***

| | | | | |
|---|---|---|---|---|
| FRET E | 0.25 | 0.25 | 0.76 | 0.76 |
| kinetic model | 0 | 0.0772712 | 0 | 0.724288 |
| | 0.0799077 | 0 | 0 | 0 |
| | 0 | 0 | 0 | 0.0818057 |
| | 0.605016 | 0 | 0.0710662 | 0 |

***6) SMACKS***

| | | | | |
|---|---|---|---|---|
| FRET E | 0.24 | 0.24 | 0.76 | 0.76 |
| kinetic model | 0 | 0 | 0.068 | 0 |
| | 0.007 | 0 | 0.622 | 0.164 |
| | 0.107 | 0.749 | 0 | 0 |
| | 0 | 0.077 | 0.017 | 0 |

##### *7) Correlation*

| | | | |
|---|---|---|---|
| FRET E | 0.22 | 0.76 | 0.9 |
| kinetic model | n.a. | n.a. | n.a. |
| | n.a. | n.a. | n.a. |
| | n.a. | n.a. | n.a. |

##### *8) Edge finding (CK)*

n.a.

##### *9) Edge finding (k-means)*

n.a.

##### *10) Step finding (2 FRET states)*

| | | |
|---|---|---|
| FRET E | 0.221 | 0.802 |
| kinetic model | 0 | 0.522 |
| | 0.669 | 0 |

##### *10b) Step finding (3 FRET states)*

| | | | |
|---|---|---|---|
| FRET E | 0.217 | 0.618 | 0.851 |
| kinetic model | 0 | 0.295 | 0.153 |
| | 0.57 | 0 | 0.256 |
| | 0.238 | 0.182 | 0 |

##### *11) STaSI*

| | | | | | | | | |
|---|---|---|---|---|---|---|---|---|
| FRET E | 0.17 | 0.25 | 0.38 | 0.54 | 0.68 | 0.76 | 0.85 | 0.92 |
| kinetic model | 0 | 0.17 | 0.085 | 0.132 | 0.076 | 0.054 | 0.033 | 0.033 |
| | 0.078 | 0 | 0.08 | 0.157 | 0.094 | 0.078 | 0.09 | 0.025 |
| | 0.294 | 0.631 | 0 | 0.381 | 0.268 | 0.251 | 0.199 | 0.078 |
| | 0.356 | 0.717 | 0.169 | 0 | 0.327 | 0.423 | 0.305 | 0.198 |
| | 0.195 | 0.33 | 0.131 | 0.207 | 0 | 0.178 | 0.246 | 0.119 |
| | 0.061 | 0.228 | 0.102 | 0.282 | 0.126 | 0 | 0.105 | 0.071 |
| | 0.054 | 0.124 | 0.07 | 0.172 | 0.175 | 0.073 | 0 | 0.059 |
| | 0.051 | 0.082 | 0.048 | 0.065 | 0.106 | 0.092 | 0.065 | 0 |

##### *12) MASH-FRET (bootstrap)*

| | | | | |
|---|---|---|---|---|
| FRET E | 0.451 | 0.863 | 0.227 | 0.702 |
| kinetic model | 0 | 0.193 | 0.843 | 0.455 |
| | 0.106 | 0 | 0.264 | 0.273 |
| | 0.176 | 0.279 | 0 | 0.221 |
| | 0.194 | 0.466 | 0.353 | 0 |

***13) MASH-FRET (probabilistic)***

| | | | | |
|---|---|---|---|---|
| FRET E | 0.251 | 0.251 | 0.743 | 0.743 |
| kinetic model | 0 | 0.015 | 0.062 | 0.677 |
| | 0 | 0 | 0 | 0.072 |
| | 0.014 | 0.003 | 0 | 0.028 |
| | 0.569 | 0.072 | 0 | 0 |

***14) postFRET (2 FRET states)***

| | | |
|---|---|---|
| FRET E | 0.24 | 0.81 |
| kinetic model | 0 | 0.2070822 |
| | 0.0711783 | 0 |

***14b) postFRET (3 FRET states)***

| | | | |
|---|---|---|---|
| FRET E | 0.23 | 0.5 | 0.8 |
| kinetic model | 0 | 0.0009144 | 0.0738384 |
| | 0.0058854 | 0 | 0.0400443 |
| | 0.0638665 | 0.0002925 | 0 |

***14c) postFRET (4 FRET states)***

| | | | | |
|---|---|---|---|---|
| FRET E | 0.25 | 0.5 | 0.69 | 0.85 |
| kinetic model | 0 | 0.0011741 | 0.08894954 | 0.10552059 |
| | 0.48463045 | 0 | 0.30084519 | 0.01306089 |
| | 0.03141345 | 0.29437991 | 0 | 0.04156456 |
| | 0.11838752 | 0.00822046 | 0.04257101 | 0 |

**Supplementary Tables 3**

Kinetic models for the data shown in Fig. 5d-f of the main text, inferred by the participating groups with the specified tools. Units of the rate constants are in $s^{-1}$. The full submission, including standard deviations of the FRET efficiencies and uncertainties of the rate constants, can be found in the Supplementary Datafiles.

***1) Pomegranate***

| | | | | |
|---|---|---|---|---|
| FRET E | 0.208 | 0.507 | 0.703 | 0.93 |
| kinetic model | 0 | 0.754 | 0.916 | 0.817 |
| | 3.101 | 0 | 2.773 | 5.424 |
| | 1.144 | 1.088 | 0 | 1.779 |
| | 0.763 | 1.133 | 0.945 | 0 |

***2) Tracy (HMM)***

| | | | |
|---|---|---|---|
| FRET E | 0.23 | 0.76 | 0.9 |
| kinetic model | 0 | 0.038 | 0 |
| | 0.042 | 0 | 0 |
| | 0.23 | 0.52 | 0 |

***3) FRETboard***

| | | | | |
|---|---|---|---|---|
| FRET E | 0.267 | 0.565 | 0.726 | 0.849 |
| kinetic model | 0 | 0.066317 | 0.180713 | 0.117159 |
| | 0.754011 | 0 | 0.545455 | 0.069519 |
| | 0.448457 | 0.121142 | 0 | 0.189866 |
| | 0.183439 | 0.049089 | 0.258365 | 0 |

***4) Hidden-Markury***

| | | |
|---|---|---|
| FRET E | 0.243 | 0.795 |
| kinetic model | 0 | 0.523 |
| | 0.492 | 0 |

***5) SMACKS(SS) (2 FRET states)***

| | | | | |
|---|---|---|---|---|
| FRET E | 0.26 | 0.26 | 0.77 | 0.77 |
| kinetic model | 0 | 0.0464469 | 0 | 0.75673 |
| | 0.0777646 | 0 | 0 | 0 |
| | 0 | 0 | 0 | 0.0767983 |
| | 0.676816 | 0 | 0.0375133 | 0 |

***5b) SMACKS(SS) (3 FRET states)***

| | | | | |
|---|---|---|---|---|
| FRET E | 0.24 | 0.62 | 0.62 | 0.81 |
| kinetic model | 0 | 0.308986 | 0.119066 | 0.162909 |
| | 1.77861 | 0 | 0 | 1.61935 |
| | 0.200454 | 0 | 0 | 0.0708802 |
| | 0.307538 | 0.613002 | 0.0852028 | 0 |

##### *6) SMACKS*

| | | | | |
|---|---|---|---|---|
| FRET E | 0.26 | 0.26 | 0.73 | 0.73 |
| kinetic model | 0 | 0 | 0.069 | 0 |
| | 0.006 | 0 | 0.668 | 0.102 |
| | 0.049 | 0.813 | 0 | 0 |
| | 0 | 0.081 | 0.041 | 0 |

##### *7) Correlation*

| | | | |
|---|---|---|---|
| FRET E | 0.23 | 0.75 | 0.88 |
| kinetic model | n.a. | n.a. | n.a. |
| | n.a. | n.a. | n.a. |
| | n.a. | n.a. | n.a. |

##### *8) Edge finding (CK)*

n.a.

##### *9) Edge finding (k-means)*

n.a.

##### *10) Step finding (2 FRET states)*

| | | |
|---|---|---|
| FRET E | 0.243 | 0.795 |
| kinetic model | 0 | 0.565 |
| | 0.588 | 0 |

##### *10b) Step finding (3 FRET states)*

| | | | |
|---|---|---|---|
| FRET E | 0.243 | 0.74 | 0.87 |
| kinetic model | 0 | 0.436 | 0.121 |
| | 0.6 | 0 | 0.067 |
| | 0.292 | 0.09 | 0 |

##### *11) STaSI*

| | | | | |
|---|---|---|---|---|
| FRET E | 0.25 | 0.25 | 0.8 | 0.8 |
| kinetic model | n.a. | n.a. | n.a. | n.a. |
| | n.a. | n.a. | n.a. | n.a. |
| | n.a. | n.a. | n.a. | n.a. |
| | n.a. | n.a. | n.a. | n.a. |

##### *12) MASH-FRET (bootstrap)*

| | | |
|---|---|---|
| FRET E | 0.755 | 0.271 |
| kinetic model | 0 | 0.53 |
| | 0.66 | 0 |

***13) MASH-FRET (probabilistic)***

| | | | | |
|---|---|---|---|---|
| FRET E | 0.27 | 0.27 | 0.75 | 0.75 |
| kinetic model | 0 | 0 | 0 | 0.043 |
| | 0.006 | 0 | 0.047 | 0.728 |
| | 0.003 | 0.062 | 0 | 0 |
| | 0.026 | 0.691 | 0 | 0 |

***14) postFRET (2 FRET states)***

| | | |
|---|---|---|
| FRET E | 0.26 | 0.8 |
| kinetic model | 0 | 0.338698 |
| | 0.345414 | 0 |

***14b) postFRET (3 FRET states)***

| | | | |
|---|---|---|---|
| FRET E | 0.25 | 0.65 | 0.85 |
| kinetic model | 0 | 0.00109842 | 0.21834065 |
| | 0.00301381 | 0 | 0.08283056 |
| | 0.27579656 | 0.00353179 | 0 |

***14c) postFRET (4 FRET states)***

| | | | | |
|---|---|---|---|---|
| FRET E | 0.25 | 0.49 | 0.69 | 0.85 |
| kinetic model | 0 | 0.0002070865 | 0.1452989654 | 0.2005696159 |
| | 0.2068609902 | 0 | 0.8709210325 | 0.0475529494 |
| | 0.4412775024 | 0.0248333492 | 0 | 0.0361363519 |
| | 0.2763561503 | 0.0829865525 | 0.006026129 | 0 |

**Supplementary Tables 4**

Kinetic models for the data shown in Fig. 5g-i of the main text, inferred by the participating groups with the specified tools. Units of the rate constants are in $s^{-1}$. The full submission, including standard deviations of the FRET efficiencies and uncertainties of the rate constants, can be found in the Supplementary Datafiles.

***1) Pomegranate***

| | | | | |
|---|---|---|---|---|
| FRET E | 0.239 | 0.479 | 0.706 | 0.897 |
| rate model | 0 | 0.504 | 0.588 | 0.755 |
| | 2.171 | 0 | 2.944 | 4.19 |
| | 0.946 | 0.612 | 0 | 1.147 |
| | 0.505 | 0.597 | 0.684 | 0 |

***2) Tracy (HMM)***

| | | | |
|---|---|---|---|
| FRET E | 0.23 | 0.76 | 0.9 |
| rate model | 0 | 0.019 | 0.017 |
| | 0.048 | 0 | 0.027 |
| | 0.011 | 0.007 | 0 |

***3) FRETboard***

| | | | | |
|---|---|---|---|---|
| FRET E | 0.257 | 0.691 | 0.806 | 0.909 |
| rate model | 0 | 0.106142 | 0.1358 | 0.045267 |
| | 0.173973 | 0 | 0.280822 | 0.078082 |
| | 0.243874 | 0.231039 | 0 | 0.038506 |
| | 0.063406 | 0.083031 | 0.076993 | 0 |

***4) Hidden-Markury***

| | | |
|---|---|---|
| FRET E | 0.237 | 0.815 |
| rate model | 0 | 0.376 |
| | 0.256 | 0 |

***5) SMACKS(SS)***

| | | | | |
|---|---|---|---|---|
| FRET E | 0.27 | 0.27 | 0.79 | 0.79 |
| rate model | 0 | 0.0181472 | 0 | 0.593785 |
| | 0.0505473 | 0 | 0 | 0 |
| | 0 | 0 | 0 | 0.270328 |
| | 0.578138 | 0 | 0.431789 | 0 |

***6) SMACKS***

| | | | | |
|---|---|---|---|---|
| FRET E | 0.27 | 0.27 | 0.78 | 0.78 |
| rate model | 0 | 0 | 0.032 | 0 |
| | 0 | 0 | 0.442 | 0.228 |
| | 0.108 | 0.926 | 0 | 0.393 |
| | 0 | 0.141 | 0 | 0 |

***7) Correlation***

| | | | |
|---|---|---|---|
| FRET E | 0.23 | 0.75 | 0.87 |
| rate model | n.a. | n.a. | n.a. |
| | n.a. | n.a. | n.a. |
| | n.a. | n.a. | n.a. |

***8) Edge finding (CK)***

n.a.

***9) Edge finding (k-means)***

n.a.

***10) Step finding (2 FRET states)***

| | | |
|---|---|---|
| FRET E | 0.237 | 0.815 |
| rate model | 0 | 0.443 |
| | 0.444 | 0 |

***10b) Step finding (3 FRET states)***

| | | | |
|---|---|---|---|
| FRET E | 0.234 | 0.722 | 0.862 |
| rate model | 0 | 0.318 | 0.106 |
| | 0.374 | 0 | 0.107 |
| | 0.126 | 0.118 | 0 |

***11) STaSI***

| | | | | | | |
|---|---|---|---|---|---|---|
| FRET E | 0.14 | 0.26 | 0.54 | 0.72 | 0.85 | 0.95 |
| rate model | 0 | 0.179 | 0.129 | 0.082 | 0.014 | 0.014 |
| | 0.039 | 0 | 0.188 | 0.198 | 0.089 | 0.025 |
| | 0.209 | 0.921 | 0 | 0.585 | 0.381 | 0.095 |
| | 0.028 | 0.298 | 0.174 | 0 | 0.241 | 0.071 |
| | 0.007 | 0.133 | 0.1 | 0.186 | 0 | 0.028 |
| | 0.01 | 0.068 | 0.06 | 0.123 | 0.053 | 0 |

***12) MASH-FRET (bootstrap)***

| | | | | | | |
|---|---|---|---|---|---|---|
| FRET E | 0.267 | 0.847 | 0.668 | 0.668 | 0.668 | 0.668 |
| rate model | 0 | 0.224 | 0.007 | 0.012 | 0.084 | 0.145 |
| | 0.166 | 0 | 0.006 | 0.01 | 0.071 | 0.123 |
| | 1.749 | 1.325 | 0 | 0 | 0 | 0 |
| | 1.749 | 0.271 | 0 | 0 | 0 | 0 |
| | 0.221 | 1.325 | 0 | 0 | 0 | 0 |
| | 0.221 | 0.271 | 0 | 0 | 0 | 0 |

***13) MASH-FRET (probabilistic)***

| | | | | |
|---|---|---|---|---|
| FRET E | 0.298 | 0.298 | 0.777 | 0.777 |
| rate model | 0 | 0 | 0.017 | 0.53 |
| | 0 | 0 | 0 | 0.009 |
| | 0.013 | 0 | 0 | 0 |
| | 0.321 | 0.003 | 0 | 0 |

***14) postFRET (2 FRET states)***

| | | |
|---|---|---|
| FRET E | 0.27 | 0.83 |
| rate model | 0 | 0.0304298 |
| | 0.0217805 | 0 |

***14b) postFRET (3 FRET states)***

| | | | |
|---|---|---|---|
| FRET E | 0.25 | 0.59 | 0.83 |
| rate model | 0 | 0.00032411 | 0.02923996 |
| | 0.05430609 | 0 | 0.07807773 |
| | 0.00175319 | 0.04096638 | 0 |

***14c) postFRET (4 FRET states)***

| | | | | |
|---|---|---|---|---|
| FRET E | 0.25 | 0.59 | 0.78 | 0.91 |
| rate model | 0 | 0.0537498259 | 0.0468467928 | 0.0352511436 |
| | 0.2059474513 | 0 | 0.5658220374 | 0.0844386522 |
| | 0.1690830952 | 0.4614106448 | 0 | 0.247915522 |
| | 0.0291453273 | 0.0784158076 | 0.0674403618 | 0 |
