## Supplementary material for "Inferring kinetic rate constants from single-molecule FRET trajectories – a blind benchmark of kinetic analysis tools": MATLAB Simulator: collapseStates.m

```
%collapse states to degenerate states

function mStateTime = collapseStates(mTransitionRate, lookupCollapse, mStateTime)
	if (size(lookupCollapse,1) ~= size(mTransitionRate,1))
		msg = [getBaseName() ' : Dimension mismatch.'];
		disp(msg);
	end
	
	
	mState = mStateTime(1:end-1,1);		%omit last entry as it is 0
	mStateClps = lookupCollapse(mState(:));
	
	mStateTime(1:end-1,2) = mStateClps;
	mStateTime(end,2) = 0;
end
```
