## Supplementary material for "Inferring kinetic rate constants from single-molecule FRET trajectories – a blind benchmark of kinetic analysis tools": MATLAB Simulator: Config_simulation_round_1_level_2.cfg

```
%parameter version 190618


%transition rate matrix
-0.680	+0.300	+0.380
+0.050	-0.420	+0.370
+0.450	+0.100	-0.550

	%transition rates (NOT transition probablities)
	%row #: inital state
	%col #: final state
	%for states and row/col number, counting starts with 1
	%for example q_{32}=+0.850 is the rate to go from state 3 to state 2
	%matrix has to be NxN (where N is number of states) and rows have to sum to zero
	%in units of 1/s


%frameRate
10
	%frame rate in units of 1/s


%bleachRate
0.02
	%bleach rate in units of 1/s


%minimum trace length
10
	%minimum trace length in seconds


%maximum trace length
200
	%maximum trace length in seconds


%lookupCollapse
1	2	3
	%vector that connects states to degenerate states
	%for example "1 1 2 2" means that states 1 and 2 are mapped to the degenerate state 1
	%and states 3 and 4 are both mapped to degenerate state 2
	%if no collapse is needed, just use 1 2 3 ... N


%include blinking
0
	%flag to include blinking of dyes in the simulation (0 don't include, 1 include)


%donor blinking kinetics
-0.03	+0.03
+15.0	-15.0
	%2x2 matrix for now, state1 is On (bright), state2 is Off (dark)
	%transition rates (NOT transition probablities)


%acceptor blinking kinetics
-0.03	+0.03
+15.0	-15.0
	%2x2 matrix for now, state1 is On (bright), state2 is Off (dark)
	%transition rates (NOT transition probablities)


%fluorescence intensity, mean
7200	800		8000	%(degenerate) state 1
3750	4250	8000	%(degenerate) state 2
800		7200	8000	%(degenerate) state 3
0		0		8000	%Ddark state
8000	0		0		%Adark state
	%cols: intensity for channels Idd, Ida and Iaa
	%rows: (degenerate) states
	%make sure Idd+Ida=const for all (degenerate) states


%fluorescence intensity, covariance matrix
0.9e6	0		0		%(degenerate) state 1
0		0.5e6	0
0		0		1.2e6

0.6e6	0		0		%(degenerate) state 2
0		0.6e6	0
0		0		1.2e6

0.5e6	0		0		%(degenerate) state 3
0		0.9e6	0
0		0		1.2e6

0.2e6	0		0		%Ddark state
0		0.2e6	0
0		0		1.2e6

1.2e6	0		0		%Adark state
0		0.2e6	0
0		0		0.2e6
	%per matrix: covariance matrix for each (degenerate) state
	%rows, cols: in the same channel order as for the mean intensity
	%if, for example, the distribution of each intensity channel
	%in each degenerate state should have a width (sigma of a normal dist) of 1000
	%and no correlation with other channels, then 1000^2 on the diagonal


%per trace level variability
5
	%parameter that controls per trace level heterogeneity
	%the intensity level for each state and intensity channel is drawn
	%from a normal distribution with mu as given in <fluorescence intensity, mean>
	%and a sigma that is sqrt(mu)*parameter
	%turn off by setting to 0


%per trace brightness variability
1.2
	%parameter that controls per trace brightness heterogeneity
	%an uniformly distributed random number r is drawn from the interval (-1, 1)
	%and all intensities of the current molecule are multiplied by n^r;
	%thus a per trace variability perameter of 2 would result in a 
	%multiplication by a factor from the interval (0.5, 2)
	%turn off by setting to 0


%number of traces
150
```
