## Supplementary material for "Inferring kinetic rate constants from single-molecule FRET trajectories – a blind benchmark of kinetic analysis tools": MATLAB Simulator: doSimContTime.m

```
%simulates a trace
%returns a matrix with state, deg. state, dwell start, dwell length

function mStateTime = doSimContTime(mTransitionRate, bleachRate, minTraceLen, maxTraceLen, verbose)
	%mTransitionRate	NumStates x NumStates; row#: inital state, col#: final state
	%					in unit of 1/s
	%					rows sum to zero
	%bleachRate			in unit of 1/s
	%minTraceLen		in units of s
	%maxTraceLen		in units of s
	%verbose			default (0): shut up
	%
	%mStateTime			[state, degenerated state, start time dwell, dwell time]
	
	
	
	%% Set default values and check input
	%in matlab matching of arguments to variables is done positionally, so there is no way to skip an argument
	%pass [] to indicate a skipped argument
	
	if (isempty(verbose))
		verbose = 0;
	end
	
	if (isempty(minTraceLen))
		minTraceLen = 0;
	end
	
	if (maxTraceLen == 0)
		maxTraceLen = [];
	end
	
	if ( isempty(bleachRate) && isempty(maxTraceLen) )
		basename = getBaseName();
		msg = [basename ': Either bleachRate or maxTraceLen need to be set'];
		error(msg)
	end
	
	
	
	%% Get start state (draw randomly accroding to stationary disttribution)
	
	%get stationary distribution
	mStationaryDist = getStationaryDist(mTransitionRate);
	
	%get random start state according to stationary distribution
	mStationaryDist_cum = cumsum(mStationaryDist);
	
	r = rand;	%rand returns random number in ]0;1[
	
	for i = 1:size(mStationaryDist, 1)
		if (mStationaryDist_cum(i) > r)
			currS = i;
			if (verbose)
				fprintf('%s : r=%f, currS=%d\n',getBaseName(),r,currS);
			end
			break
		end
	end
	
	
	
	%% Get trace length from exponential distribution
	if ~isempty(maxTraceLen)
		traceLength = maxTraceLen;
	end
	
	if ~isempty(bleachRate)
		while 1
			r = rand;
			traceLength = -1/bleachRate*log(1-r);	%log(X) returns the natural logarithm ln(x) of each element in array X
			if (traceLength > minTraceLen)
				break
			end
		end
	end
	
	%check if traceLength exceeds maxTraceLen
	if ~isempty(maxTraceLen)
		if (traceLength > maxTraceLen)
			traceLength = maxTraceLen;
		end
	end
	
	if (verbose)
		fprintf('%s : traceLength=%f\n',getBaseName(),traceLength);
	end
	
	
	
	%% Simulate the dwells
	j = 1;
	mStateTime(1,1) = currS;
	mStateTime(1,3) = 0;
	
	while 1
		shortestDwell = +inf;
		%get dwell time for each depopulating transition
		for i =1:size(mStationaryDist, 1)
			if (i == currS)		%skip state that is currently populated
				continue
			end
			transRate = mTransitionRate(currS,i);
			r = rand;
			dwell = -1/transRate*log(1-r);	%log(X) returns the natural logarithm ln(x) of each element in array X
			
			if (dwell<shortestDwell)		%shortest dwell wins
				GoToState = i;
				shortestDwell = dwell;
			end
		end
		
		mStateTime(j,4) = shortestDwell;
		mStateTime(j+1,1) = GoToState;
		mStateTime(j+1,3) = mStateTime(j,3) + shortestDwell;
		
		if (verbose)
			fprintf('%s : GoToState=%d, shortestDwell=%f\n',getBaseName(),GoToState,shortestDwell);
		end
		
		if (mStateTime(j+1,3) > traceLength)	%bleached
			%fprintf('j=%d, traceLength=%f, shortestDwell=%f\n', j, traceLength, shortestDwell);
			mStateTime(j,4) = traceLength - mStateTime(j,3);
			mStateTime(j+1,1) = 0;
			mStateTime(j+1,3) = traceLength;
			mStateTime(j+1,4) = +inf;
			break
		else
			j = j + 1;
			currS = GoToState;
		end
	end
	
	if (verbose)
		%hFig = figure;
		stairs(mStateTime(:,3), mStateTime(:,1:2));
	end
end
```
