## Supplementary material for "Inferring kinetic rate constants from single-molecule FRET trajectories – a blind benchmark of kinetic analysis tools": MATLAB Simulator: getBlinkTrajectory.m

```
function mStateTime = getBlinkTrajectory(mBlinkRate, traceLength)
%mBlinkRate			2x2 matrix for now, state1 is On (bright), state2 is Off (dark)
%					in units of 1/s
%					rows sum to zero
%traceLength		length of the simulated trace
%					in units of s


verbose = 0;

%get stationary distribution
mStationaryDist = getStationaryDist(mBlinkRate);


%get random start state according to stationary distribution
mStationaryDist_cum = cumsum(mStationaryDist);

r = rand;	%rand returns random number in ]0;1[

for i = 1:size(mStationaryDist, 1)
	if (mStationaryDist_cum(i) > r)
		currState = i;
		if (verbose)
			fprintf('%s : r=%f, currS=%d\n',getBaseName(),r,currState);
		end
		break
	end
end


j = 1;
mStateTime(1,1) = currState;
mStateTime(1,3) = 0;

while 1
	shortestDwell = +inf;
	%get dwell time for each depopulating transition
	for i =1:size(mStationaryDist, 1)
		if (i == currState)		%skip state that is currently populated
			continue
		end
		transRate = mBlinkRate(currState,i);
		r = rand;
		dwell = -1/transRate*log(1-r);	%log(X) returns the natural logarithm ln(x) of each element in array X
		
		if (dwell<shortestDwell)		%shortest dwell wins
			nextState = i;
			shortestDwell = dwell;
		end
	end
	
	mStateTime(j,4) = shortestDwell;
	mStateTime(j+1,1) = nextState;
	mStateTime(j+1,3) = mStateTime(j,3) + shortestDwell;
	
	if (verbose)
		fprintf('%s : GoToState=%d, shortestDwell=%f\n',getBaseName(),nextState,shortestDwell);
	end
	
	if (mStateTime(j+1,3) > traceLength)	%bleached
		%fprintf('j=%d, traceLength=%f, shortestDwell=%f\n', j, traceLength, shortestDwell);
		mStateTime(j,4) = traceLength - mStateTime(j,3);
		mStateTime(j+1,1) = 0;
		mStateTime(j+1,3) = traceLength;
		mStateTime(j+1,4) = +inf;
		break
	else
		j = j + 1;
		currState = nextState;
	end
end

%copy state trajectory from row 1 to row 2
mStateTime(:,2) = mStateTime(:,1);


if (verbose)
	%hFig = figure;
	stairs(mStateTime(:,3), mStateTime(:,1:2));
end
end
```
