## Supplementary material for "Inferring kinetic rate constants from single-molecule FRET trajectories – a blind benchmark of kinetic analysis tools": MATLAB Simulator: getRatioStateBlink.m

```
function ratioStateBlink = getRatioStateBlink(mStateTime_blink, tStartFrame, tEndFrame, mStateTime, numDegStates)
%ratioState			fraction of time different states are occupied
%mStateTime_blink	mSateTime for blinking, [state, 0, start time dwell, dwell time]
%tStartFrame		start time of current frame
%tEndFrame			end time of current frame
%mStateTime			mStateTime for the states [state, deg. state, start time dwell, dwell time]
%numDegStates		number of deg. states
%blinkState			1: Ddark, 2: Adark


normalize = 0;	%don't normalize ratioState to frame time
verbose = 0;


%% get a list of blink events in this frame

%mStartEnd: [state, t_start, t_end]
mStartEnd(:,1:2) = mStateTime_blink(:,[1,3]);	%omit col2
mStartEnd(:,3) = mStateTime_blink(:,3) + mStateTime_blink(:,4);

%replace the dwells with the frame start or end time
idx1 = find(mStartEnd(:,2)>tStartFrame,1);	%empty if start > traceLength
mStartEnd(idx1-1,2) = tStartFrame;
idx2 = find(mStartEnd(:,3)>tEndFrame,1);
mStartEnd(idx2,3) = tEndFrame;
mStartEnd_short = mStartEnd(idx1-1:idx2,:);
idx = (mStartEnd_short(:,1) == 2);
mStartEnd_short2 = mStartEnd_short(idx,2:3);


%% sum the time each state was populated during this blink event

ratioStateBlink = 0;
if (~isempty(mStartEnd_short2))
	for j = 1:size(mStartEnd_short2, 1)
		ratioState_tmp = getStateRatio(mStartEnd_short2(j,1), mStartEnd_short2(j,2), mStateTime, numDegStates, normalize, verbose);
		ratioStateBlink = ratioStateBlink + ratioState_tmp;
	end
end


end
```
