## Supplementary material for "Inferring kinetic rate constants from single-molecule FRET trajectories – a blind benchmark of kinetic analysis tools": MATLAB Simulator: getStateRatio2.m

```
%returns a vector of the deg. state, sampled at <samples> points, in the
%current frame

function ratioState2 = getStateRatio2(ratioState2, tStartFrame, tEndFrame, mStateTime, process)
%tStart				time point to start
%tEnd				time point to stop evaluating time interval
%mStateTime			[state, deg. state, start time dwell, dwell time]
%samples			how many sub-samples should be drawn for each frame
%process			which independend processes (here:
%					conformation biomolecule, donor blink, acc blink)


samples = size(ratioState2, 1);

for i = 1:samples		%last element of range is also included in MATLAB
	frameTime = tEndFrame - tStartFrame;
	currT = tStartFrame + (i-1)*frameTime/samples;
	
	%find corresponding row in mStartEnd
	k = find(mStateTime(:,3)>currT,1);	%empty if currT > bleach event
	if (isempty(k))		%bleached already
		ratioState2(i, process) = 0;
	else
		ratioState2(i, process) = mStateTime(k-1, 2);	%get deg. state at currT
	end


end
```
