## Supplementary material for "Inferring kinetic rate constants from single-molecule FRET trajectories – a blind benchmark of kinetic analysis tools": MATLAB Simulator: getStateRatio.m

```
function ratioState = getStateRatio(tStart, tEnd, mStateTime, numDegStates, normalize, verbose)
%tStart				time point to start
%tEnd				time point to stop evaluating time interval
%mStateTime			[state, deg. state, start time dwell, dwell time]
%numDegStates		number of degenerated states
%normalize			if 1, normalize the ratio to tEnd-tStart
%verbose			flag to be verbose or not

if (tStart < 0)
	fprintf('%s : tStart=%d has to be >= 0\n', getBaseName(), tStart);
end

if (tEnd < tStart)
	fprintf('%s : tEnd samller than tStart\n', getBaseName());
	error('Mid script reached, this error message was made on purpose')
end

if ~exist('verbose','var')
	verbose = 0;
end


%mStartEnd: [deg. state, t_start, t_end]
mStartEnd(:,1:2) = mStateTime(:,2:3);
mStartEnd(:,3) = mStateTime(:,3) + mStateTime(:,4);

%for historical reasons, rename...
start = tStart;
stop = tEnd;
frameTime = tEnd - tStart;

%find starting row in mStartEnd
k = find(mStartEnd(:,2)>start,1);	%empty if start > traceLength
j = k-1;

if isempty(k)
	disp('start not found');
end

ratioState = zeros(numDegStates,1);

while (1)
	if (j == size(mStateTime,1)-1)	%last row is only bleached
		currStart = max([start, mStartEnd(j,2)]);
		currStop = min([stop, mStartEnd(j,3)]);
		ratioState(mStartEnd(j,1)) = ratioState(mStartEnd(j,1)) + currStop - currStart;
		break
	elseif (stop<mStartEnd(j,3))
		currStart = max([start, mStartEnd(j,2)]);
		ratioState(mStartEnd(j,1)) = ratioState(mStartEnd(j,1)) + stop - currStart;
		break
	else
		currStart = max([start, mStartEnd(j,2)]);
		%this is just the dwell time in most cases, but not sure if it
		%would be easier to use the dwell time and have another if in
		%case the currStart is not the start of the dwell
		ratioState(mStartEnd(j,1)) = ratioState(mStartEnd(j,1)) + mStartEnd(j,3) - currStart;
		j = j + 1;
	end
end

if (verbose)
	fprintf('%s : frameTime=%d, ratioState (not normalized)\n',getBaseName(),frameTime);
	disp(ratioState);
end

if (normalize)
	ratioState = ratioState / frameTime;
end
end
```
