## Supplementary material for "Inferring kinetic rate constants from single-molecule FRET trajectories – a blind benchmark of kinetic analysis tools": MATLAB Simulator: getStationaryDist.m

```
%returns the stationary distribution of a transition RATE matrix
%i.e. elements of matrix are rates and rows of the matrix sum to zero
function mStationaryDist = getStationaryDist(mTransitionRate)
%not sure why the matrix has to be transformed here, but otherwise
%eigenvalues and eigenvectors are wrong
[V,D] = eig(mTransitionRate');
vEigenvalues = diag(D);

%decomposition using floating-point computations, thus result is close to, but not exactly, 0
vEigenvaluesReal = round(1e10*real(vEigenvalues))/1e10;

%test for == 0 since transition RATE matrix, if transition matrix, this would be == 1
I = find(vEigenvaluesReal == 0);

if (size(I,1) == 0)
	msg = [getBaseName() ' : Found no stationary distribution. Check input transition rate matrix.'];
	disp(msg);
	mStationaryDist = NaN;
	return
elseif (size(I,1) > 1)
	msg = [getBaseName() ' : Found more than one stationary distribution. Check input transition rate matrix.'];
	disp(msg);
	mStationaryDist = NaN;
	return
end

%check if imag(vEigenvalues) == 0
if (isreal(vEigenvalues(I(1))) == 0)
	msg = [getBaseName() ' : Found no stationary distribution. Check input transition rate matrix.'];
	disp(msg);
	mStationaryDist = NaN;
	return
end

mStationaryDist = V(:,I(1));
%normalize to sum=1
s = sum(mStationaryDist);
mStationaryDist = mStationaryDist/s;
end
```
