## Supplementary material for "Inferring kinetic rate constants from single-molecule FRET trajectories – a blind benchmark of kinetic analysis tools": MATLAB Simulator: getTrace.m

```
%Simulation of one trace
%dimensions=3 (I^d_d, I^d_a, I^a_a)

%mu x=1, y=dimensions, z=states
%sigma x=y=dimensions, z=states

%intensity x=state, y=dimensions
%mu+sigma*randn


%function [trace, mStateTime, mStateTime_blinkD, mStateTime_blinkA] = getTrace(param, R, verbose)
function structTrace = getTrace(param, R, verbose)
	transitionRateMatrix = param.transitionRateMatrix;
	bleachRate = param.bleachRate;
	minTrLen = param.minTraceLen;
	maxTrLen = param.maxTraceLen;
	lookupCollapse = param.lookupCollapse;
	mu = param.mu;
	numDims = param.numDims;
	numDegStates = param.numDegStates;
	frameRate = param.frameRate;
	rmLastFrame = param.rmLastFrame;
	trLevelVar = param.perTraceLevelVar;
	trBrightVar = param.perTraceBrightVar;
	includeBlinking = param.includeBlinking;
	mRate_blinkD = param.mRate_blinkD;
	mRate_blinkA = param.mRate_blinkA;
	
	
	%get a trace-specific multiplier for the intensities in all
	%dimensions/channels
	if (trLevelVar)
		%replace all mu with a random mu drawn from a normal distribution
		%around mu, with width sqrt(mu)*trLevelVar
		mu_sqrt = mu.^0.5;
		mu_real = normrnd(mu, mu_sqrt*trLevelVar);
		%make sure Iaa is the same for all states
		mu_real(1,3,1:end-1) = mu_real(1,3,1);
	else
		mu_real = mu;
	end
	
	
	%get continues time trace
	mStateTime = doSimContTime(transitionRateMatrix, bleachRate, minTrLen, maxTrLen, verbose);
	
	
	%collapse states that share the same observation
	mStateTime = collapseStates(transitionRateMatrix, lookupCollapse, mStateTime);
	
	
	%get blinking mStateTime; 1 is on, 2 is off
	traceLength = mStateTime(end, 3);
	if (includeBlinking)
		mStateTime_blinkD = getBlinkTrajectory(mRate_blinkD, traceLength);
		mStateTime_blinkA = getBlinkTrajectory(mRate_blinkA, traceLength);
	else
		mStateTime_blinkD = [1 0 0 traceLength; 0 0 traceLength +inf];
		mStateTime_blinkA = [1 0 0 traceLength; 0 0 traceLength +inf];
	end
	
	
	%discretize
	traceFrameMax = ceil(traceLength*frameRate);
	if (rmLastFrame)
		traceFrameMax = traceFrameMax -1;
	end
	
	trace = zeros(traceFrameMax, numDims);
	
	for i = 1:traceFrameMax
		%get sub-samples in the current frame
		samples = 100;	%each frame is sub-samples by this factor
		frameTime = 1/frameRate;
		tStartFrame = (i-1)*frameTime;
		tEndFrame = i*frameTime;
		ratioState2 = zeros(samples, 4);
		%4 rows for
		%	conformation of biomolecule
		%	overall emission state
		%	donor blink
		%	acc blink
		ratioState2 = getStateRatio2(ratioState2, tStartFrame, tEndFrame, mStateTime, 1);
		
		
		%update ratioState according to blinking
		ratioState2 = getStateRatio2(ratioState2, tStartFrame, tEndFrame, mStateTime_blinkD, 3);
		ratioState2 = getStateRatio2(ratioState2, tStartFrame, tEndFrame, mStateTime_blinkA, 4);
		
		
		ratioState2(:,2) = ratioState2(:,1);
		%Ddark and Adark (no emission at all)
		idx = (ratioState2(:,3)==2 & ratioState2(:,4)==2);
		ratioState2(idx,2) = 0;
		%Ddark
		idx = (ratioState2(:,3)==2 & ratioState2(:,4)==1);
		ratioState2(idx,2) = numDegStates+1;
		%Adark
		idx = (ratioState2(:,3)==1 & ratioState2(:,4)==2);
		ratioState2(idx,2) = numDegStates+2;
		
		ratioState = zeros(numDegStates+2, 1);
		for j = 1:(numDegStates+2)
			ratioState(j) = sum(ratioState2(:,2)==j);
		end
		
		%get random intensities for all states in this frame
		%+2 to include Ddark and Adark sates
		Z = zeros(numDegStates+2,numDims);
		for j = 1:(numDegStates+2)
			Z(j,:) = mu_real(:,:,j) + randn(1,numDims)*R(:,:,j);
		end
		
		%sum intensities according to the ratio of the corresponding state
		ratioState = ratioState / samples;
		ZZ = ratioState' * Z;
		
		%write intensities to trace
		trace(i,2:4) = ZZ;
	end
	
	%multiply intensities by random factor
	if (trBrightVar)
		%generate uniformly distributed random number in the interval (-1,1)
		r = -1 + 2*rand; %rand returns a single uniformly distributed random number in the interval (0,1)
		trBrightMultiplier = trBrightVar^r;
		trace(:,2:4) = trace(:,2:4) * trBrightMultiplier;
		%fprintf('multiplier %f\n', multiplier);
	else
		trBrightMultiplier = NaN;
	end
	
	%calculate timestamp of frame and FRET efficiency
	trace(:,1) = linspace(0,1/frameRate*(traceFrameMax-1),traceFrameMax);
	trace(:,5) = trace(:,3) ./ (trace(:,2)+trace(:,3));
	trace(:,6) = (trace(:,2)+trace(:,3)) ./ (trace(:,2)+trace(:,3)+trace(:,4));
	
	%trace(:,1)	timestamp
	%trace(:,2)	Idd
	%trace(:,3)	Ida
	%trace(:,4)	Iaa
	%trace(:,5)	FRET eff
	%trace(:,6)	stoichiometry
	
	%collect return values
	structTrace = struct;
	structTrace.Trace = trace;
	structTrace.mStateTime = mStateTime;
	structTrace.mStateTime_blinkD = mStateTime_blinkD;
	structTrace.mStateTime_blinkA = mStateTime_blinkA;
	structTrace.mu_real = mu_real;
	structTrace.trBrightMultiplier = trBrightMultiplier;
end
```
