## Supplementary material for "Inferring kinetic rate constants from single-molecule FRET trajectories – a blind benchmark of kinetic analysis tools": MATLAB Simulator: importSettings.m

```
%import parameters for the simulation from a text file

function [param, success] = importSettings(param)
	%param			struct with at least field numDims
	%				(i.e. number of recorded fluorescence channels)
	%				the current function expects numDims==3 with
	%				dimLabels = {'Idd'; 'Ida'; 'Iaa'}
	
	success = 1;
	numDims = param.numDims;
	
	[file,path] = uigetfile('*','Select the simulation config...');
	
	if isequal(file,0)		%user selected Cancel
		disp('User pressed cancel');
		success = -1;
		return
	else
		fileName = fullfile(path,file);
		param.configFileName = fileName;
	end
	
	strFileSetting = fileread(fileName);
	TextAsCells = regexp(strFileSetting, '\n', 'split');
	
	
	
	%% transition rate matrix
	idx = find(strcmp(TextAsCells, '%transition rate matrix'), 1, 'first');
	if (isempty(idx))
		disp('Could not find header ''%transition rate matrix''')
		success = -1;
		return
	end
	
	A = importdata(fileName,'\t',idx);
	if (~isfield(A, 'data'))
		A = importdata(fileName,'\t',idx+1);
		if (~isfield(A, 'data'))
			disp('No numeric data found after header ''%transition rate matrix''')
			success = -1;
			return
		end
	end
	
	numStates = size(A.data,2);	%does not stop when finding text, so crop later
	param.numStates = numStates;
	param.transitionRateMatrix = A.data(1:numStates,1:numStates);
	
	
	
	%% frame rate
	key   = '%frameRate';
	idx = strfind(strFileSetting, key);
	if (isempty(idx))
		disp('Could not find header ''%frameRate''')
		success = -1;
		return
	else
		Value = sscanf(strFileSetting(idx(1) + length(key):end), '%f');
		param.frameRate = Value;
	end
	
	
	
	%% bleach rate
	key   = '%bleachRate';
	idx = strfind(strFileSetting, key);
	if (isempty(idx))
		disp('Could not find header ''%bleachRate''')
		success = -1;
		return
	else
		Value = sscanf(strFileSetting(idx(1) + length(key):end), '%f');
		param.bleachRate = Value;
	end
	
	
	
	%% minimum trace length
	key   = '%minimum trace length';
	idx = strfind(strFileSetting, key);
	if (isempty(idx))
		disp(['Could not find header >>' key '<<'])
		success = -1;
		return
	else
		Value = sscanf(strFileSetting(idx(1) + length(key):end), '%f');
		param.minTraceLen = Value;
	end
	
	
	
	%% maximum trace length
	key   = '%maximum trace length';
	idx = strfind(strFileSetting, key);
	if (isempty(idx))
		disp(['Could not find header >>' key '<<'])
		success = -1;
		return
	else
		Value = sscanf(strFileSetting(idx(1) + length(key):end), '%f');
		param.maxTraceLen = Value;
	end
	
	
	
	%% look-up table for collapsing states to degenerate states
	key   = '%lookupCollapse';
	idx = strfind(strFileSetting, key);
	if (isempty(idx))
		disp(['Could not find header >>' key '<<'])
		success = -1;
		return
	else
		Value = sscanf(strFileSetting(idx(1) + length(key):end), '%f');
		lookupCollapse = Value;
	end
	
	if (numStates ~= size(lookupCollapse, 1))
		disp('lookupCollapse does not match the number of states given in the transition rate matrix.')
		success = -1;
		return
	else
		param.lookupCollapse = lookupCollapse;
		numDegStates = size(unique(lookupCollapse),1);	%is used further down
		param.numDegStates = numDegStates;
	end
	
	
	
	%% include blinking
	key   = '%include blinking';
	idx = strfind(strFileSetting, key);
	if (isempty(idx))
		disp(['Could not find header >>' key '<<'])
		success = -1;
		return
	else
		Value = sscanf(strFileSetting(idx(1) + length(key):end), '%f');
		param.includeBlinking = Value;
	end
	
	
	
	%% blinking kinetics donor
	key   = '%donor blinking kinetics';
	idx = find(strcmp(TextAsCells, key), 1, 'first');
	if (isempty(idx))
		disp(['Could not find header >>' key '<<'])
		success = -1;
		return
	end
		
	A = importdata(fileName,'\t',idx);
	if (~isfield(A, 'data'))
		disp(['No numeric data found after header >>' key '<<'])
		success = -1;
		return
	end
	
	len = size(A.data,2);	%does not stop when finding text, so crop later
	param.mRate_blinkD = A.data(1:len,1:len);
	
	
	
	%% blinking kinetics acceptor
	key   = '%acceptor blinking kinetics';
	idx = find(strcmp(TextAsCells, key), 1, 'first');
	if (isempty(idx))
		disp(['Could not find header >>' key '<<'])
		success = -1;
		return
	end
		
	A = importdata(fileName,'\t',idx);
	if (~isfield(A, 'data'))
		disp(['No numeric data found after header >>' key '<<'])
		success = -1;
		return
	end
	
	len = size(A.data,2);	%does not stop when finding text, so crop later
	param.mRate_blinkA = A.data(1:len,1:len);
	
	
		
	%% mean fl. intensity deg. states
	key   = '%fluorescence intensity, mean';
	idx = find(strcmp(TextAsCells, key), 1, 'first');
	if (isempty(idx))
		disp(['Could not find header >>' key '<<'])
		success = -1;
		return
	end
	
	param.mu = zeros(1,numDims,numDegStates+2);
		%not really needed, for readability, if removed, change "mu(:,:,1)=" to "mu="
		%numDegStates+2 to include Ddark and Adark states
	for i = 1:numDegStates+2
		Value = sscanf(TextAsCells{1,idx+i}, '%f', [1 3]);
		if (isempty(Value))
			disp(['Too few parameters for>>' key '<<'])
			success = -1;
			return
		else
			param.mu(:,:,i) = Value;
		end
	end
	
	
	
	%% width fl. intensity deg. states
	key   = '%fluorescence intensity, covariance matrix';
	idx = find(strcmp(TextAsCells, key), 1, 'first');
	if (isempty(idx))
		disp(['Could not find header >>' key '<<'])
		success = -1;
		return
	end
	
	param.covar = zeros(numDims,numDims,numDegStates+2);
		%not really needed, for readability, if removed, change "mu(:,:,1)=" to "mu="
		%numDegStates+2 to include Ddark and Adark states
	for i = 1:numDegStates+2
		curr = idx + (i-1)*(numDims+1);
		for j = 1:numDims
			Value = sscanf(TextAsCells{1,curr+j}, '%f', [1 3]);
			if (isempty(Value))
				disp(['Too few parameters for>>' key '<<'])
				success = -1;
				return
			else
				param.covar(j,:,i) = Value;
			end
		end
	end
	
	
	
	%% per trace level variability
	key   = '%per trace level variability';
	idx = strfind(strFileSetting, key);
	if (isempty(idx))
		disp(['Could not find header >>' key '<<'])
		success = -1;
		return
	else
		Value = sscanf(strFileSetting(idx(1) + length(key):end), '%f');
		param.perTraceLevelVar = Value;
	end
	
	
	
	%% per trace brightness variability
	key   = '%per trace brightness variability';
	idx = strfind(strFileSetting, key);
	if (isempty(idx))
		disp(['Could not find header >>' key '<<'])
		success = -1;
		return
	else
		Value = sscanf(strFileSetting(idx(1) + length(key):end), '%f');
		param.perTraceBrightVar = Value;
	end
	
	
	
	%% number of traces
	key   = '%number of traces';
	idx = strfind(strFileSetting, key);
	if (isempty(idx))
		disp('Could not find header ''%number of traces''')
		success = -1;
		return
	else
		Value = sscanf(strFileSetting(idx(1) + length(key):end), '%f');
		param.numTraces = Value;
	end

end
```
