## Supplementary material for "Inferring kinetic rate constants from single-molecule FRET trajectories – a blind benchmark of kinetic analysis tools": MATLAB Simulator: README.pdf

### Simulation of smFRET trajectories for the kinSoft Challenge

#### Installation guide & system requirements

The simulation scripts need a working installation of Matlab. No further installation steps are required.

The simulation was tested with Matlab R2011b on Windows 10, Matlab R2017b on Ubuntu 16.04.4LTS and Matlab R2019b on Pop!_OS 19.10.

#### Running simulations

1. Start Matlab.
2. Add the folder `MATLAB_simulator` to the Matlab path (navigate to the location where the `MATLAB_simulator` folder is located, then right-click > Add to Path > Selected Folders and Subfolders).
3. Open `MATLAB_simulator/simContTime.m`.
4. Run the script (by clicking the "Run" button or pressing F5).
5. A dialog will open in order for you to select a configuration file for the simulation (an example configuration file `simulation_example.cfg` is available in the folder `MATLAB_simulator`).

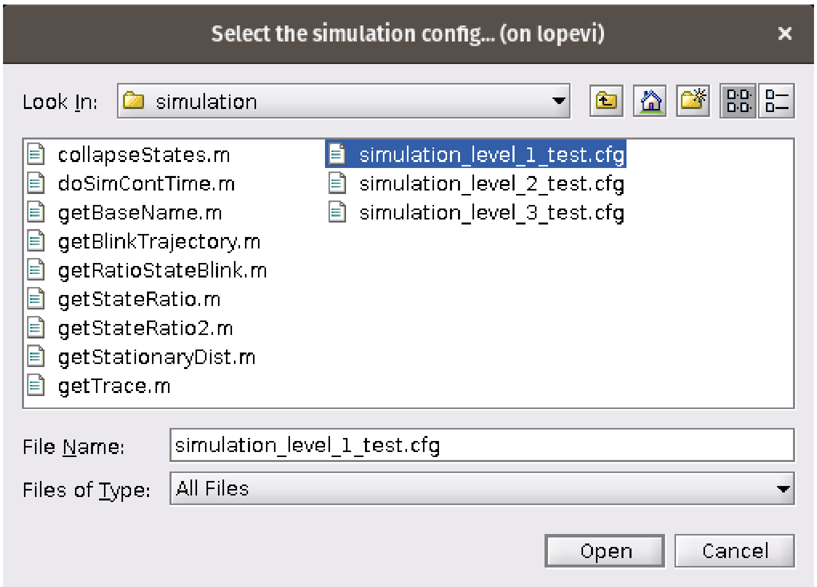


6. The simulation typically takes a few seconds, depending on the size of the kinetic model and the number of molecules to simulate.
7. A new folder called `sim_<date>_<time>` will be created in the current folder, containing:
    1. `allData.mat`: a MAT-file with the simulation parameters and all the simulated data
    2. `dwellTimes_state_<i>.txt`: the list of dwell times for state i

3. `state_time_<j>.txt`: a tab-separated list of state, degenerated state, t_start (s), and t_dwell (s) for molecule j
4. `trace_<j>.txt`: a tab-separated list of time, fluorescence intensity and FRET efficiency for molecule j

### Re-creating the challenge datasets

To exactly re-create the challenge datasets, two settings are needed: First, the seed for the random number generated (RNG) that was used during the initial simulation. Second, the configuration file for the simulation. Both can be found in the folder of the corresponding challenge dataset.

The RNG seed is stored in `allData.mat` (variable name `param.rngSeed`). Use this integer as seed in line 55 of `simContTime.m` after uncommenting. For convenience, the seeds for the three challenge datasets are also given below:

| Dataset | RNG seed |
|---|---|
| level 1 | 1046468070 |
| level 2 | 1256105342 |
| level 3 | 2593404440 |

Finally, run the script as detailed above and select the configuration file used for this challenge dataset.
