## Supplementary material for "Inferring kinetic rate constants from single-molecule FRET trajectories – a blind benchmark of kinetic analysis tools": MATLAB Simulator: simContTime.m

```
%% Version 190629

%% ToDo
% * to get cont time with bleaching
%   ** slide time along all 3 mStateTime to merge them
%   ** then old version of getStateRatio() should work again
% * blinking of D and A could be included in one mTransRate
% * sanity check for neg. values on diagonal and pos values else in trans rate matrix


%% Tested With
%(R2011b on Windows 10)
%R2017b on Ubuntu 16.04.4 LTS


%% How It Works
%discretization:
%for each frame and for each intensity channel
%	get the ratio of how long each state is populated during this frame
%	(this is done by subsampling by a factor of 100)
%	draw a random intensity from a gaussian distribution for each state
%	sum intensities, weight according to ratios of each state


%% Fields in struct param
%param.rngSeed
%param.dimLabels (hardcoded)
%param.numDims (hardcoded)
%param.rmLastFrame (hardcoded)
%param.configFileName
%param.numStates
%param.transitionRateMatrix
%param.frameRate
%param.bleachRate
%param.minTraceLen
%param.maxTraceLen
%param.lookupCollapse
%param.numDegStates
%param.includeBlinking
%param.mRate_blinkD
%param.mRate_blinkA
%param.mu
%param.covar
%param.perTraceLevelVar
%param.perTraceBrightVar
%param.numTraces


function simContTime()

%% Preamble

clear all
seed = randi(2^32-1);
%seed = 3792279088; disp('Hardcoded seed')
%rng('shuffle')  %seeds the random number generator based on the current time
rng(seed)

param.dimLabels = {'Idd'; 'Ida'; 'Iaa'};	%actually not used anywhere
param.numDims = size(param.dimLabels, 1);
param.rmLastFrame = 1;
param.rngSeed = seed;
verbose = 0;


%% Import

[param, success] = importSettings(param);

tic

if (success ~= 1)
	disp('Import failed')
	return
end


%Cholesky factorization, formula from help on randn
%sqrt(covar) on main diagonal
%numDegStates+2 to include Ddark and Adark
%R = zeros(param.numDims,param.numDims,param.numDegStates);
R = zeros(param.numDims,param.numDims,param.numDegStates+2);
%for i = 1:param.numDegStates
for i = 1:param.numDegStates+2
	R(:,:,i) = chol(param.covar(:,:,i));
end


%% Sanity Check import

%check minTraceLen vs bleachRate
if (param.minTraceLen*10 > 1/param.bleachRate)
	disp('The supplied bleach rate is large compared to the minimal trace length.');
end

%check if rows in trans rate matrix sum to zero
testSum = sum(param.transitionRateMatrix, 2);
testSum = round(1e10*testSum)/1e10;
F = find(testSum ~= 0);
if (size(F, 1) > 0)
	g = sprintf('%d ', F);
	fprintf('The supplied transition rate matrix does not sum to zero in row(s): %s\n', g);
	disp('Aborting.');
	return
end

%check that mu_DA + mu_DD = const
testSum = param.mu(1,1,1:end-2) + param.mu(1,2,1:end-2);	%end-2 to exclude Ddark, Adark
testSum = testSum(:);	%reshape into a single column vector
testSum = testSum - testSum(1);
testSum = sum(testSum, 1);
if (testSum ~= 0)
	disp('The supplied mean fluorescence intensities for Idd+Ida are not constant for all states.');
	disp('Aborting.');
	return
end

%check that per trace brightness variability parameter is within reasonable bounds
trVar = param.perTraceBrightVar;
if (trVar < 0 || trVar > 5)
	disp('The supplied per trace variablity parameter is not within reasonable bounds.');
	disp('Aborting.');
	return
end


%% Simulation of all traces

cellAllTraces(param.numTraces,1:6) = {[]};
cellDwellTimes(param.numStates,1) = {[]};

for i = 1:param.numTraces
%	structTrace = struct([]);
	structCurrTrace = getTrace(param, R, verbose);
	currTrace = structCurrTrace.Trace;
	mStateTime = structCurrTrace.mStateTime;
	mStateTime_blinkD = structCurrTrace.mStateTime_blinkD;
	mStateTime_blinkA = structCurrTrace.mStateTime_blinkA;
	mu_real = structCurrTrace.mu_real;
	trBrightMultiplier = structCurrTrace.trBrightMultiplier;
	
	cellAllTraces(i,:) = {currTrace; mStateTime; mStateTime_blinkD; mStateTime_blinkA; mu_real; trBrightMultiplier};
	
	for j = 1:(size(mStateTime,1)-1)		%omit last dwell, as it is inf
		currState = mStateTime(j,1);
		cellDwellTimes(currState) = {[cellDwellTimes{currState}; mStateTime(j,4)]};
	end
end


%% Export

folderNameBase = 'sim_';
formatOut = 'yymmdd_HHMMSS';
folderNameDate = datestr(now,formatOut);

folderName = [folderNameBase folderNameDate];
[status,msg] = mkdir(folderName);

if ~status
	disp('Could not create a folder in the current directory.')
	disp(msg)
	return
end

saveCDF = pwd;
cd(folderName);

%loop over all traces and save them
fileNameTrace = 'trace_';
fileNameStateTime = 'state_time_';

for i = 1:param.numTraces
	currTrace = cellAllTraces{i,1};
	currTrace = currTrace(:,1:5);	%omit column 6, which holds stoichiometry
	fileID = fopen([fileNameTrace num2str(i) '.txt'], 'w');
	fprintf(fileID, '%%t (s)\tIdd (a.u.)\tIda (a.u.)\tIaa (a.u.)\tFRET E\n');
	fprintf(fileID, '%.3f\t%+.3e\t%+.3e\t%+.3e\t%+.3f\n', currTrace.');
	fclose(fileID);
	
	currStateTime = cellAllTraces{i,2};
	fileID = fopen([fileNameStateTime num2str(i) '.txt'], 'w');
	fprintf(fileID, '%%state\tdeg. state\tt_start (s)\tt_dwell (s)\n');
	fprintf(fileID, '%d\t%d\t%+.3e\t%+.3e\n', currStateTime.');
	fclose(fileID);
end


%loop over all states and save cellDwellTimes
fileNameDT = 'dwellTimes_state_';
for i = 1:param.numStates
	currDT = cellDwellTimes{i,1};
	fileID = fopen([fileNameDT num2str(i) '.txt'], 'w');
	fprintf(fileID, ['%%dwell times of state ' num2str(i) ' (s)\n']);
	fprintf(fileID, '%.6f\n', currDT.');
	fclose(fileID);
end


%save config file
[~,name,ext] = fileparts(param.configFileName);
copyfile(param.configFileName, ['Copy_of_' name ext]);


%write variable to allData.mat
save('allData.mat', 'cellAllTraces', 'cellDwellTimes', 'param');


%change back to original working dir
cd(saveCDF);


%% Write out variables to workspace

assignin('base','cellAllTraces',cellAllTraces);
assignin('base','cellDwellTimes',cellDwellTimes);
assignin('base','param',param);


%% Print out some useful info
toc

mStationaryDist = getStationaryDist(param.transitionRateMatrix);
disp(mStationaryDist)

end


%% Tests for differents parts of the code

function testMatlabRand()
	r = rand(100);	%100x100 matrix of uniformly distributed random numbers
	r_range = [min(r(:)) max(r(:))]
end


function testGetStationaryDist(mTransitionRate)
	len = 1e7;	%1e10 to get close to input
	verbose = 0;
	mStationaryDist = getStationaryDist(mTransitionRate);
	
	occState = zeros(size(mStationaryDist,1),1);
	
	%get random start state according to stationary distribution
	mStationaryDist_cum = cumsum(mStationaryDist);
	
	for i = 1:len
	
		r = rand;	%rand returns random number in ]0;1[
		
		for i = 1:size(mStationaryDist, 1)
			if (mStationaryDist_cum(i) > r)
				currS = i;
				if (verbose)
					fprintf('%s : r=%f, currS=%d\n',getBaseName(),r,currS);
				end
				break
			end
		end
		
		occState(currS) = occState(currS) + 1;

	end
	
	mStationaryDist
	occState = occState / len
	
end


function [counts,centers] = testDistTraceLength()
	len = 1e5;
	traceLen = zeros(len, 1);
	bleachRate = 0.02;	%in unit of 1/time, roughly 0.02 for Hsp90
	
	
	for i = 1:len
		%get trace length from exponential distribution
		r = rand;
		traceLength = -1/bleachRate*log(1-r);	%log(X) returns the natural logarithm ln(x) of each element in array X
		
		traceLen(i) = traceLength;
	end
	
	[counts,centers] = hist(traceLen);
	bar(centers,counts);
	
	%[counts,centers] = testDistTraceLength()
end


function [z, counts,centers] = testMultivariatGauss()

	%from help on randn
	%but watch out, sigma should actually be covariance matrix
	%mu = [1 2];
	%sigma = [1 0.5; 0.5 2];
	%R = chol(sigma);
	%z = repmat(mu,10,1) + randn(10,2)*R
	
	mu = [2 10];
	%sigma = [1 0.5; 0.5 2];
	sigma = [1 0; 0 2];
	R = chol(sigma);
	z = repmat(mu,1e4,1) + randn(1e4,2)*R;
	
	[counts,centers] = hist(z(:,2),20);
	
	%matlab gaussian function returns c1 that is sqrt(2) too large
	%difference between normal dist and matlab funtion
	
	
	
%[z, counts,centers] = testMultivariatGauss();

%%figure;
%%scatter(mR(:,1), mR(:,2));
%%linkdata on
end
```
