## Supplementary figures and images for "Inferring kinetic rate constants from single-molecule FRET trajectories – a blind benchmark of kinetic analysis tools"

### Screenshot from 2021-11-18 10-32-39.png

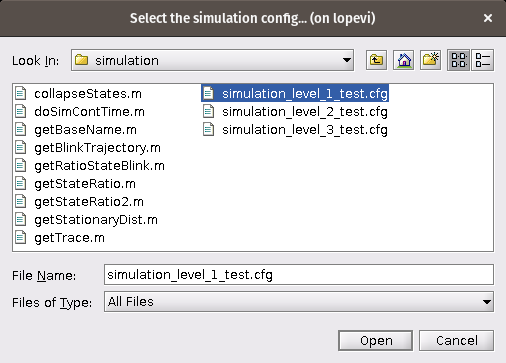
